# Larval environment reshapes mosquito disease risk via phenotypic and molecular plasticity

**DOI:** 10.1101/2025.06.16.659972

**Authors:** Karthikeyan Chandrasegaran, Melody Walker, Jeffrey M. Marano, Spruha Rami, Adaline Bisese, James Weger-Lucarelli, Chloé Lahondère, Michael A. Robert, Lauren M. Childs, Clément Vinauger

**Affiliations:** Department of Biochemistry, Virginia Polytechnic Institute and State University, Blacksburg, VA 24061, USA; Department of Mathematics, Virginia Polytechnic Institute and State University, Blacksburg, VA 24061, USA; Department of Biomedical Sciences and Pathobiology, VA-MD College of Veterinary Medicine, Blacksburg, VA 24061, USA; Fralin Life Sciences Institute, Virginia Polytechnic Institute and State University, Blacksburg, VA 24061, USA; Center for Emerging Zoonotic and Arthropod-borne Pathogens, Virginia Polytechnic Institute and State University, Blacksburg, VA 24061, USA; Center for the Mathematics of Biosystems, Virginia Polytechnic Institute and State University, Blacksburg, VA 24061, USA; Department of Entomology, University of California Riverside, Riverside, CA 92521, USA; Center for Pharmacometrics and Systems Pharmacology, Department of Pharmaceutics, College of Pharmacy, University of Florida, Orlando, FL 328227, USA; Department of Biomedical Sciences, College of Veterinary Medicine and Biomedical Sciences, Colorado State University, Colorado, USA

**Keywords:** Transstadial Effects, Epidemiology, Mosquito Neuroecology, Size Dependency, Zika, Mathematical Model

## Abstract

Early-life environmental conditions can exert profound, lasting effects on adult phenotypes, with major consequences for fitness and disease transmission, especially in holometabolous insects like mosquitoes, which are a key vector species. Yet, the molecular mechanisms through which juvenile environments shape adult physiology and behavior via transstadial effects remain largely unresolved. Here, we demonstrate that larval competition, a key ecological stressor, profoundly alters adult body size, survival, reproductive output, host-seeking behavior, olfactory neurophysiology, and vector competence in the mosquito *Aedes aegypti*. Crucially, using transcriptomic profiling and integrative network analyses, we identify seven regulatory hub genes whose expression is strongly associated with size-dependent variation in olfactory behavior, reproductive investment, and Zika virus transmission potential. These hub genes belong to gene modules enriched for functions in chemosensory processing, metabolic regulation, and signal transduction, revealing a molecular framework mediating environmentally induced plasticity across metamorphosis. Integrating these empirical findings into a transmission model, we show that incomplete larval control can inadvertently increase outbreak risk by producing larger, longer-lived, and more competent vectors. Our results uncover molecular mechanisms underpinning phenotypic plasticity in disease vectors and highlight the critical need to account for transstadial effects in models of vector-borne disease transmission.

## Introduction

Mosquito-borne diseases such as malaria, Zika, dengue, and chikungunya continue to pose severe global health challenges. With projections indicating that nearly half of the world’s population could be at risk by 2050 (1) and annual death tolls exceeding one million (2), these diseases demand urgent, coordinated intervention. Efforts to control their spread are increasingly challenged by rapid urbanization, the emergence of insecticide resistance, the global dispersal of invasive mosquito species, and the accelerating effects of climate change (3). These factors collectively reshape the ecological conditions mosquitoes experience during their development, with potentially profound yet poorly understood consequences for adult vectorial capacity and disease transmission dynamics.

While it is well established that environmental conditions during the aquatic juvenile stages of mosquitoes influence adult phenotypes, including lifespan, fecundity, and pathogen susceptibility (4, 5), the mechanistic pathways through which these early-life effects persist across complete metamorphosis remain largely unresolved. This knowledge gap limits our ability to predict how environmental variability and anthropogenic change will shape disease risk at both individual and population scales. In particular, although correlations between larval environment, adult body size, and transmission potential have been noted, it remains unknown how growing conditions shape adult mosquito physiology and behavior through coordinated molecular and neural mechanisms.

Here, we address this critical gap by combining ecologically relevant manipulations of larval competition in *Aedes aegypti* with comprehensive phenotypic assays, olfactory neurophysiology, transcriptomic profiling, arbovirus transmission assays, and mathematical modeling. We show that larval density profoundly impacts adult mosquitoes, not only altering their size and reproductive output but also modulating host-seeking behavior, responses to repellents, olfactory integration, and Zika virus transmission potential. Through integrative transcriptomic analyses, we identify seven regulatory hub genes that mediate size-dependent variation in these traits, uncovering a molecular framework by which early environmental stress leaves persistent effects on adult phenotypes. Notably, we demonstrate that these effects extend to the neural representation of host-seeking cues, where carbon dioxide (CO_2_) and host volatiles bidirectionally modulate one another’s encoding in the mosquito brain — challenging the conventional view of CO_2_ as a simple activator of host-seeking.

Finally, by integrating these fine-scale mechanistic findings into a dynamic disease transmission model, we show that incomplete larval control strategies may unintentionally amplify outbreak risk by favoring the emergence of larger, longer-lived, and more competent vectors. Together, our results reveal the molecular and neural pathways mediating transstadial plasticity in a key disease vector, providing new insight into how environmental variability shapes vector competence and disease dynamics, with important implications for public health interventions under conditions of climate and land-use change.

## Results and Discussion

### Growing conditions shape adult body size, life history traits, and their relationships

Mosquito growing conditions vary drastically as a function of the oviposition sites adult females select to lay their eggs (6, 7). Predators, nutrient availability, and environmental properties determine larval development time and adult size at emergence (8–11). For example, crowded larval conditions typically result in extended development time, reduced larval survival rates, and small-sized adult mosquitoes due to resource competition (9, 11). Conversely, less crowded conditions tend to produce larger-sized individuals with shorter development times, as resources per capita are more abundant (12, 13). However, the influence of larval growing conditions on adults’ behavioral response to host cues, an important contributor to the contact rate between mosquitoes and their hosts, has been overlooked. Furthermore, body size appears as a factor that covaries with multiple traits (10, 11, 14, 15). Notably, these relationships remain to be quantified and leveraged in predictive models. To bridge this knowledge gap, we examined how size variations driven by growing conditions (i.e., transstadial effects) influence the life history, host-seeking behavior, sensory physiology, and Zika virus transmission potential of *Ae. aegypti* females.

To simulate ideal and suboptimal larval growing conditions that are ecologically relevant in the laboratory (16), we reared mosquitoes under two density conditions (Fig. 1A, high and low) while keeping all other environmental parameters constant. Larval growing conditions significantly influenced mosquitoes’ development time (Fig. 1B) and survival to adulthood (Fig. 1C), both quantified using a high-throughput automated system (Fig. 1A). Females, but not males, in the high-density cohorts developed slower than those from low-density cohorts (Fig. 1B; ANOVA: *F*_1,449_ = 10.62, *p* < 0.01), likely due to heightened resource competition, creating suboptimal growth conditions (17, 18). Females emerged later than males across both cohorts (Fig. 1B; ANOVA: *F*_1,449_ =114.70, *p* < 0.001), a known pattern of protandry in mosquitoes, where males typically mature faster to maximize early mating opportunities. High-density conditions also reduced the proportion of individuals surviving to adulthood (Fig. 1C; Binomial GLM: *β* = 0.667, *z* = 3.41, *p* < 0.001). However, sex ratios remained stable across cohorts (Fig. 1D; ANOVA: *F*_1,29_ = 0.31, *p* > 0.05). High larval density not only slowed development but also produced smaller adults (lower body mass), likely due to a reduced accumulation of energy reserves (Fig. 1E). In contrast, larvae from low-density cohorts emerged as larger individuals (Fig. 1E; ANOVA: *F*_1,246_ = 588.02, *p* < 0.001). Body mass and wing length were significantly correlated, and we used either metric to measure body size, consistent with previously published studies (Fig. 1F; Linear regression: *R* = 0.83, *p* < 0.001) (11, 19). We thus used these metrics across a series of experiments to characterize the life history traits of females emerging from low and high larval density conditions, which, from this point on and for simplicity, we will refer to as large and small females, respectively. Large females, having accumulated greater energy reserves, survived significantly longer than their small-sized counterparts when provided *ad libitum* access to only DI water (Fig. 1G; Cox proportional hazards model: *β* = -1.12, HR = 0.33, *p* < 0.001). However, when provided *ad libitum* access to 10% sucrose, this survival advantage was no longer observable (Fig. 1H; Cox proportional hazards model: *β* = 0.24, HR = 1.27, *p* > 0.05), as sugar feeding allowed small females to replenish their energy reserves. This difference in longevity between sugar-fed and water-fed mosquitoes is consistent with previous observations (20) and underscores the critical role of sugar-feeding in adult mosquito survival.

**Fig. 1.**
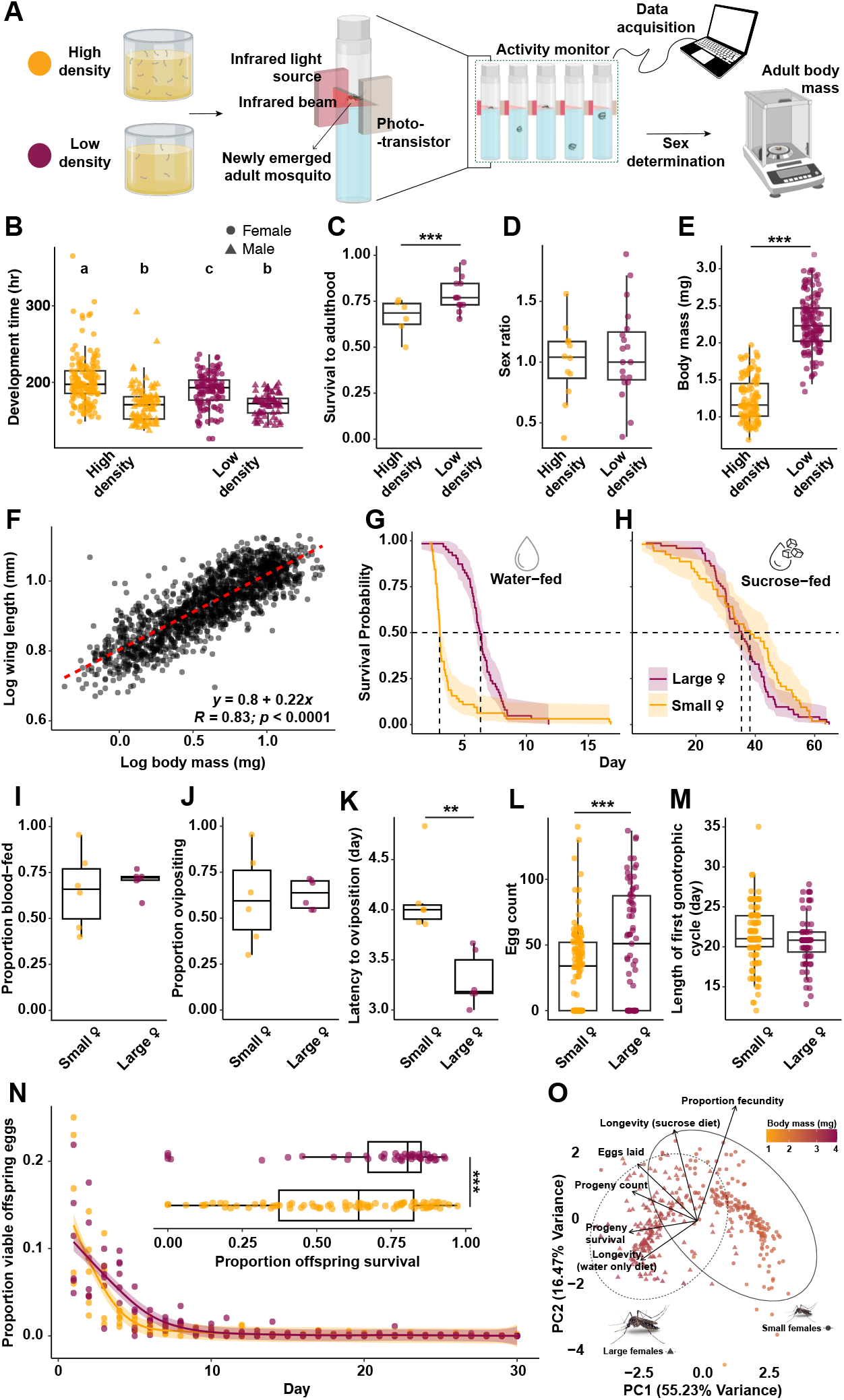
Larval growing conditions shape mosquito life history. **(A)** Schematic of the experimental approach. *Left* : Mosquito larvae were reared under either high or low density conditions. *Right* : Upon pupation, mosquitoes were transferred into glass tubes bisected by an infrared beam positioned above the water surface, which was used to detect and record the day and time of emergence. Adults were weighted within 3-6 hours post emergence. **(B)** Development time from egg hatch to adult emergence (in hours) for males (triangles) and females (circles) from high-density (orange) and low-density (maroon) conditions (high density: males, n=160 from N=7 replicates; females, n=103 from N=4 replicates; low density: males, n=119 from N=13 replicates; females, n=71 from N=7 replicates; Generalized linear model, *p* < 0.05). **(C)** Proportion of female larvae surviving to adulthood from high- and low-density conditions (high density: N=6; low density: N=13). **(D)** Sex ratio at emergence for high- and low-density conditions (high density: N=12; low density: N=19). **(E)** Adult body mass (mg) at emergence for females from high- and low-density conditions (n=115 for maroon, 133 for orange). **(F)** Log-transformed correlation between adult body mass (mg) and wing length (mm) across all mosquitoes used in this study (n=1571), accounting for allometric scaling. **(G)** Survival probability of small (orange, n=64) and large (maroon, n=63) females reared under high- and low-density conditions, respectively, provided with DI water only or **(H)** ad libitum 10% sucrose (n=53 and n=73, respectively). Dashed lines indicate the time at which survival probability reached 0.5. **(I)** Proportion of females that successfully ingested a blood meal when presented with a membrane feeder on days 6-11 post-emergence (high density: N=6; Low density: N=6; Generalized linear model, *p* > 0.05). **(J)** Proportion of blood-fed females that oviposited within 30 days post-blood meal (high density: N=6; Low density: N=6; Generalized linear model, *p* > 0.05). **(K)** Latency to oviposition, measured as the number of days from blood feeding to the first egg being laid (high density: N=6; Low density: N=6; ANOVA,*p* < 0.05). **(L)** Total eggs laid over 30 days post blood meal (high density: n=133 from N=6 replicates; low density: n=71 from N=6 replicates; Zero-Inflated Negative Binomial model, *p* < 0.05). **(M)** Length of the first gonotrophic cycle, measured as the number of days from adult emergence to the last egg laid (Linear mixed-effects model, *p* > 0.05). **(N)** Proportion of eggs from large (marron) and small females (orange) that hatched over a 30-day period, with the overall survival to the fourth larval instar shown in the inset. **(O)** PCA summarizing all quantified traits in a multidimensional space. Throughout the figure, large females and their progeny are color-coded in maroon and small females in orange. Boxplots indicate the median (horizontal black line), first and third quartiles (box limits), and 95% confidence intervals (vertical lines). Asterisks denote statistical significance (*, *p* < 0.05; **, *p* < 0.01; ***, *p* < 0.001). Letters denote significant differences for multi-factor comparisons. Exact sample sizes and summary statistics for each assay are provided as source data.

Growing conditions also significantly influenced mosquitoes’ reproductive behaviors. While similar proportions of females blood-fed (Fig. 1I; Binomial GLM: *β* = 0.23, *z* = 0.73, *p* > 0.05) and oviposited (Fig. 1J; Binomial GLM: *β* = 0.14, *z* = 0.45, *p* > 0.05), larger females laid eggs earlier post-blood feeding (Fig. 1K; Linear regression: *t* = -4.35, *p* < 0.01) and laid significantly more eggs (Fig. 1L; Zero-inflated negative binomial GLM: *β* = 0.47, *t* = 3.37, *p* < 0.001). However, the proportion of viable eggs (Zero-inflated negative binomial GLM: *β* = 0.24, *t* = 1.72, *p* > 0.05) and the length of the first gonotrophic cycle—from adult emergence to laying the last egg—were comparable between large and small females (Fig. 1M; Linear mixed-effects model: *β* = -0.76, *t* = -1.59, *p* > 0.05). Similarly, the daily proportion of viable eggs over a 30-day period did not differ between size classes (Fig. 1N; Linear mixed-effects model: *β* = 0.0025, *t* = 1.31, *p* > 0.05) but the overall proportion of offspring that survived till the last larval instar was significantly higher for large females than for smaller females (Fig. 11N inset; Linear mixed-effects model: *β* = 0.095, *t* = 3.82, *p* < 0.001). Altogether, large mosquitoes not only started laying eggs earlier and produced more eggs, but their offspring are also of higher quality, with greater chances of survival, further underscoring the fitness advantages conferred by larger body size.

Females of varying body sizes occupy distinct regions of the multivariate trait space (Principal Component Analysis, Fig. 1O), with the largest females clustering separately from their smallest counterparts, reflecting greater reproductive output of larger females (along PC1, explaining 55.23% of the variance) and differences in survival and reproductive efficiency (along PC2, 16.47% of the variance). These results suggest that larger females convert blood meals more efficiently into viable offspring with access to sugar feeding. Importantly, we observed an overlap between the largest females emerging from high larval density conditions and the smallest females from low-density conditions, indicating that differences are truly mediated by body size and not confounded by the specifics of our experimental treatments. Life history traits are central parameters for population dynamics and epidemiological modeling and, overall, our findings demonstrate the multifaceted fitness advantages of a larger body size.

### Mosquitoes’ host and plant seeking preferences are body-size dependent

Contact rates between mosquitoes and their hosts are also shaped by mosquitoes’ behavioral responses to host cues, particularly olfactory ones (14, 21–25). In this context, we subjected 5-7-day-old mosquitoes to Y-maze olfactometer assays to test whether larval density could influence the olfactory behavior of large and small mosquitoes. Specifically, we established simplified artificial blends to avoid confounding effects due to inter- and intraindividual variations in odor profiles, and quantified behavioral responses to host and plant odors, as well as CO_2_, an important host cue for host seeking (Fig. 2), and analyzed the behavioral preferences of body size bins sufficient to provide high statistical power (>0.85; n=43) and covering the entire range of body sizes sampled in our assays (see methods for details). Small females from the treatment group exhibited no significant preference for the host artificial blend (Fig. 2A1) and showed no significant preference between a plant and a host blend (Fig. 2B1) but demonstrated a significant attraction for CO_2_ (Fig. 2C1). However, this attraction was lost when CO_2_ was combined with the host odor blend (Fig. 2D1). In contrast, large females from the treatment group showed a significant preference for the host odor blend (Fig. 2A2), CO_2_ (Fig. 2C2), and the combination of CO_2_ and host odor blend (Fig. 2D2) but were equally attracted to the host and the plant odor blends (Fig. 2B2). These findings indicate that while the response to CO_2_ alone is consistent across size classes, the valence of host and plant odorants is size-dependent, regardless of the presence or absence of CO_2_.

**Fig. 2.**
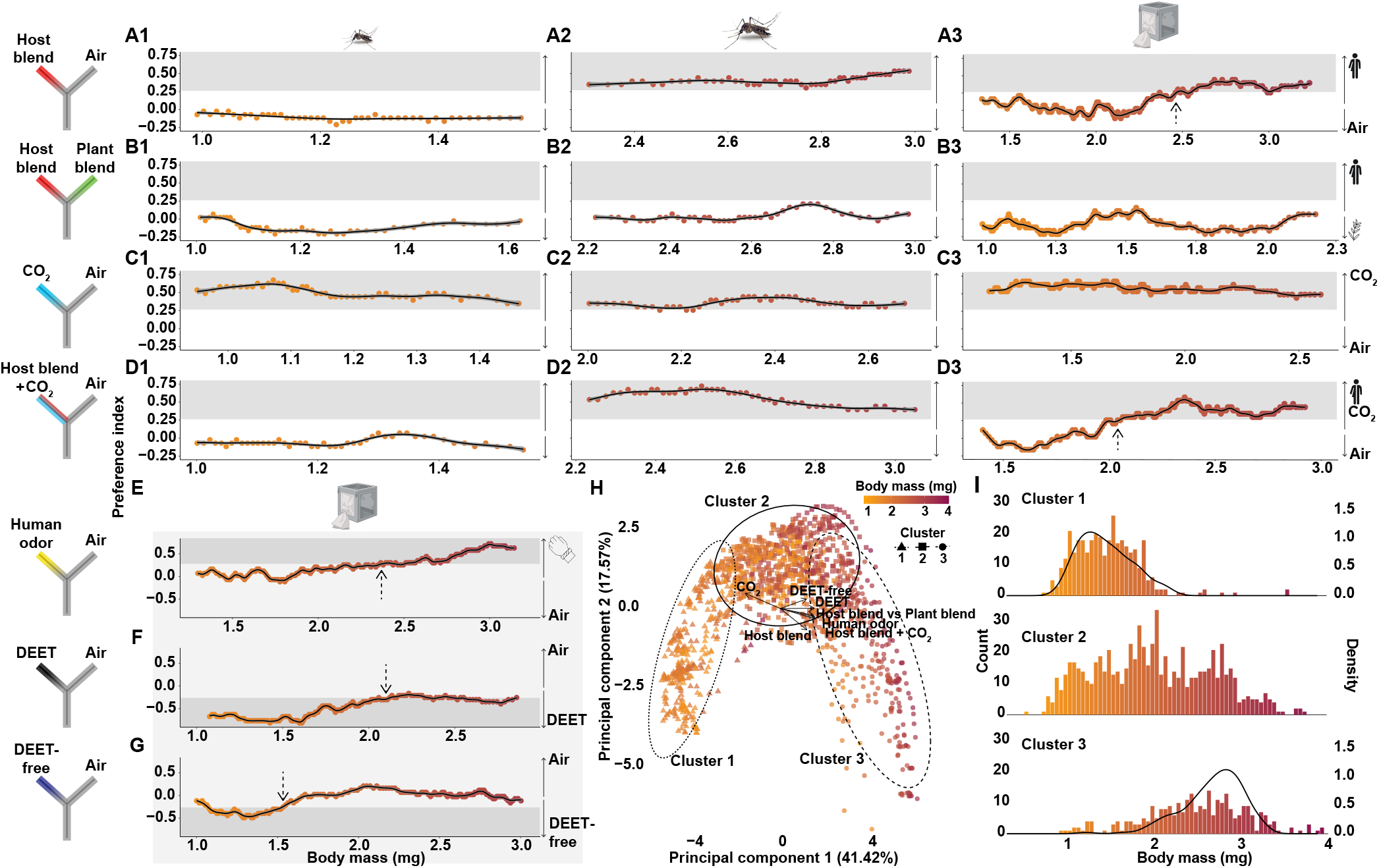
Mosquitoes’ olfactory responses are body size-dependent. Mosquito olfactory preferences to odorant stimuli were assessed in a Y-maze olfactometer. **(A–D)** Preference indices of small females reared at high density **(A1–D1)**, large females reared at low density **(A2–D2)**, and colony-derived females spanning the entire body size distribution **(A3–D3)**. Stimuli included host blend vs. air **(A)**, host vs. plant blend **(B)**, CO_2_ vs. air **(C)**, and host blend + CO_2_ vs. air **(D). (E)** Attraction to human volunteers samples assessed against a clean air control. **(F–G)** Repellent avoidance was tested by presenting DEET **(F)** or a DEET-free repellent **(G)** against clean air control. **(H)** PCA summarizing body size-dependent variation in olfactory responses across all assays (A3, B3, C3, D3, E-G). **(I)** Body size distributions of the three clusters identified in **(H)**. The preference index represents the proportion of mosquitoes selecting the odor stimulus minus the proportion choosing the control, ranging from -1 (complete avoidance) to +1 (complete preference). Data points overlapping with the shaded gray regions indicate a response significantly different from a random distribution of 50% in each arm of the olfactometer, determined using a one-tailed Binomial Exact test. The trend line of each plot was computed using a Generalized Additive Model. Exact sample sizes and summary statistics are provided as source data.

Since effects on physiological traits highly co-vary with body size, in addition to mosquitoes from our low- and high-density treatments, we also tested colony-derived individuals to evaluate whether the behavioral effects of growing conditions are also truly body size-dependent and independent of the specific experimental treatments used in our study. For this, groups of mosquitoes were raised under standard laboratory conditions (*sensu* MR4 guidelines (26)) without any deliberate attempt at controlling the amount of resources per capita (27–29). The behavioral responses of colony-derived females mirrored those of small and large females, reinforcing that host-seeking behavior is size-dependent (Fig. 2A3-D3). The maximum response to the combined stimuli (Preference index = 0.58 ± 0.06) was greater than to host odor alone (Preference index = 0.44 *±* 0.07), suggesting a potential synergistic effect between host odors and CO_2_. In addition, a body mass threshold of 2.06 mg was identified for significant preference in assays involving both the host odor blend and CO_2_ (Fig. 2D3), compared to 2.47 mg for the host odor blend alone (Fig. 2A3).

To evaluate whether similar effects would apply to more complex blends of host volatiles, we tested colony-derived mosquitoes’ responses to a nylon sleeve worn by a human volunteer (VT IRB protocol # 20-037) and found that body size also influenced responses to complex human scent (Fig. 2E). The inflection point for host-seeking behavior in these assays (2.37 mg) was within the range of inflection points observed in experiments using the artificial host blend, though with a stronger maximum response magnitude (Preference index = 0.72 ± 0.05).

Next, we evaluated the impact of body size on the behavioral responses to commercially available repellents. Colony-derived females were tested against DEET (98.11% DEET, Repel 100 Insect repellent, Bridgeton, MO, USA) and a natural repellent (30% oil of lemon eucalyptus, Repel Plant-Based Lemon Eucalyptus Insect Repellent, Bridgeton, MO, USA) (Fig. 2F,G). Smaller females showed significant avoidance of both repellents, with DEET being more effective and producing a stronger response magnitude (2F,G). Females weighing under 2.06 mg significantly avoided DEET, while avoidance of the natural repellent was observed only in mosquitoes weighing below 1.52 mg. While larger females showed a near-significant trend of avoiding DEET (Fig. 2F), they were indifferent to the natural repellent (Fig. 2G).

To better characterize the behavioral profile of adult females of varied body masses, we integrated the responses of mosquitoes in each size bin to each of the stimuli we tested in a multivariate space (PCA; Fig. 2H). In this behavioral trait space, mosquitoes distributed along a continuous arc of increasing body mass. Notably, an unsupervised clustering analysis (K means) identified 3 clusters (Fig. 2I) whose mass distribution overlaps with the distribution of masses in our small (cluster 1) and large (cluster 2) treatment groups (Fig. 1E), and centered around the median mass of our colony-derived mosquitoes (cluster 3). Together, these results provide the first demonstration that olfactory behaviors, including host-seeking and repellent avoidance, are influenced by mosquito body size.

### Mosquitoes’ olfactory sensitivity to host and plant odors is body-size independent

A possible mechanistic explanation for the size-dependent responses to odorants is that small and large females differ in their olfactory sensitivity (30), more specifically, in the tuning of their sensitivity to specific odorants since large females were more attracted to host volatiles but small females showed a greater aversion to repellents (Fig. 2). To test this hypothesis, we performed electroantennogram (EAGs) recordings on 6 day old small and large females from the two larval density treatments. 90% (n=18 out of 20) of high density (small) and 77.8% (n=14 out of 18) of low density (large) females exhibited a detectable response to a high concentration of benzaldehyde (10:100 volume:volume dilution, i.e., 1.82 × 10^*−*3^ M), used as a positive control (Binomial exact test: probability = 0.563, *p* > 0.05). Only preparations that showed a detectable response to the positive control were included in comparisons of small and large female responses (Fig. S3 A-D). Among them, no significant differences were observed in the proportion of preparations responsive to the panel of host and plant-related odorants (Fig. S3B, Binomial exact test: *p* > 0.05). However, large females exhibited higher absolute responses to stimuli from the panel (Fig. S3C; ANOVA: F_10,330_ = 1.86, *p* < 0.05), possibly due to their larger antennae size. In addition, even after normalizing responses to the response magnitude to the positive control, large females still showed a significantly greater response magnitude than small females (Fig. 3A,B; ANOVA: F_1,30_ = 7.14, *p* < 0.05). While the greater olfactory sensitivity of large females could explain the size-dependent response to host cues, this would not explain the stronger avoidance of repellents by small females. Instead, these behavioral results suggest a size-mediated modulation of the olfactory tuning to specific odorants. Next, we thus normalized responses to stimuli to that of the positive control. In this case, we observed no significant differences in responses to host and plant-related odorants (Fig. 3C; ANOVA: F_10,330_ =0.73, *p* > 0.05). In other words, the difference in olfactory preference of small and large mosquitoes cannot be fully explained by a divergence in odorant-specific sensitivity.

**Fig. 3.**
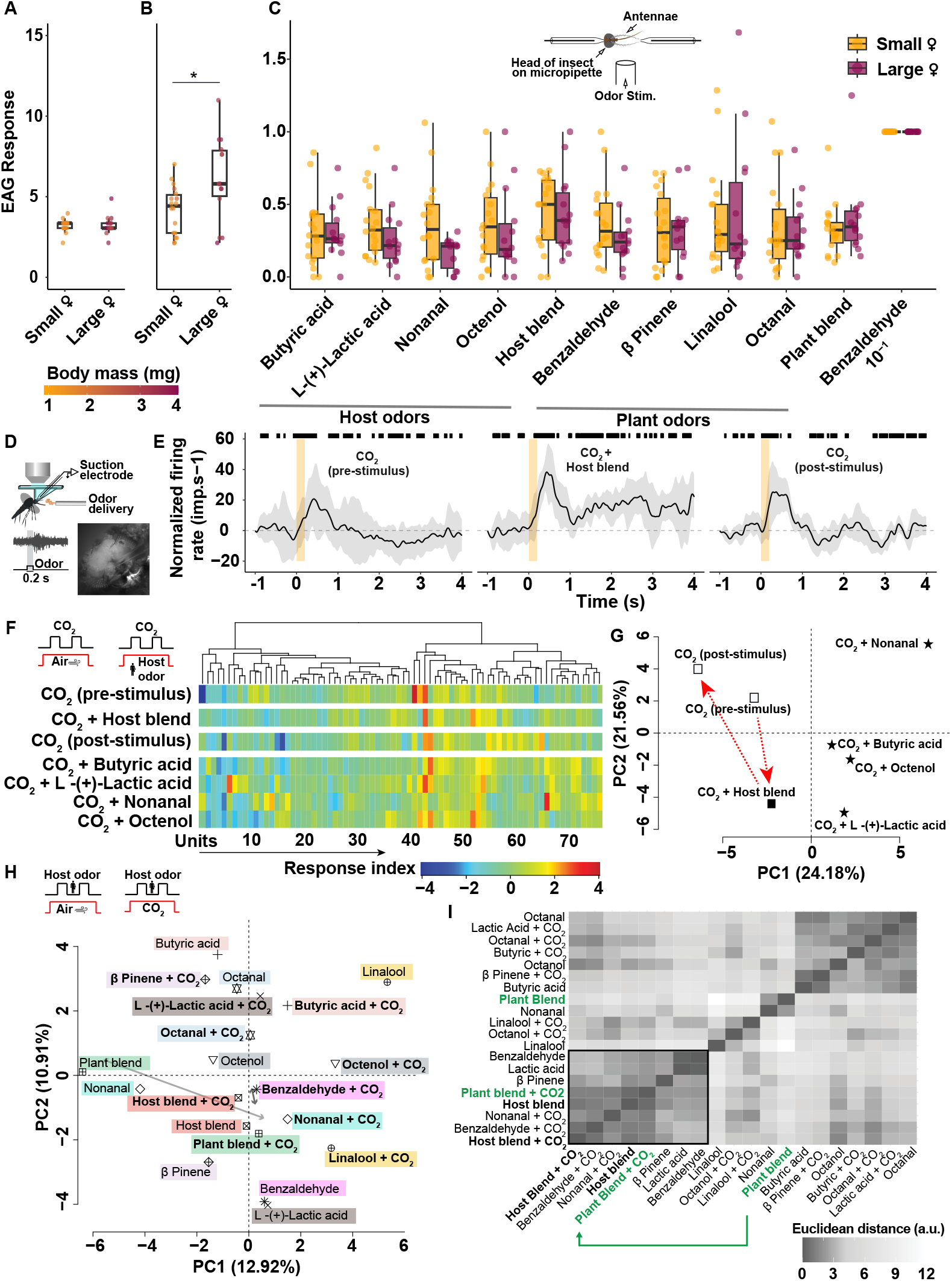
Olfactory sensitivity is body size-independent, but CO_2_ modulates encoding of host and plant odorants in the mosquito brain. **(A–C)** Electroantennogram (EAG) responses of small and large females to odorant stimuli. **(A)** Response to mineral oil (negative control). **(B)** Response to benzaldehyde (positive control, 10^*−*1^ concentration) normalized to the negative control. **(C)** Responses to the host odor blend and its constituents (butyric acid, L-(+)-lactic acid, nonanal, and octenol), the plant odor blend and its constituents (benzaldehyde, *β*-pinene, linalool, and octanal), normalized to the positive control (10^*−*1^ concentration). Each data point represents an individual mosquito’s response. (C inset) Schematic of the electrophysiological setup for EAG recordings. Statistical significance was determined using a two-factor ANOVA with Tukey post-hoc test and Bonferroni correction (*p* < 0.05). **(D–I)** Extracellular neural recordings from the mosquito antennal lobe (AL). **(D)** Schematic of the suction electrode setup, showing neural input from the antenna and maxillary palps. A representative raw trace depicts neural responses to a 200 ms pulse of host odor blend. **(E)** Raster plots and peri-event histograms display the mean (*±* SE) responses of a neural unit to CO_2_ (pre-stimulus), CO_2_ + host blend, and CO_2_ (post-stimulus) pulses, indicated by vertical orange bars. **(F)** Neural ensemble response matrix depicting the activity of 75 neural units across 13 preparations in response to CO_2_ alone, CO_2_ + host blend, and CO_2_ with individual host blend constituents. **(G)** PCA of the neural ensemble responses. CO_2_ -alone responses (both pre- and post-stimulus) and responses to CO_2_ combined with the host blend are depicted as open and closed squares, respectively. Responses to CO_2_ with individual host odor constituents are indicated as stars. The red arrow denotes the distinct shift in neural representation of CO_2_ in a host blend background. **(H)** PCA of neural responses to host and plant odors, with and without CO_2_. Each shape and color represents an odor pulse. **(I)** Distance matrix depicting the Euclidean distance between responses to a given pair of odor stimuli, with or without CO_2_. Highlighted in green, the plant blend, which is initially distinct from the host blend, shifts closer to the host odor representation in the presence of CO_2_. Exact sample sizes and statistical analyses are provided in the source data.

### Bidirectional interaction between carbon dioxide and odorants neural encoding

In our behavioral assays, the response to the combination of CO_2_ and host volatiles was body-size dependent. However, body size influenced the response to host volatiles presented alone, but not to CO_2_. This suggests a central modulation of the CO_2_ representation by host volatiles (Fig. 2A,C). To test this hypothesis, we first stimulated female mosquitoes with pulses of CO_2_ either alone or in combination with host volatile organic compounds (VOCs). Using *in vivo* extracellular recordings from the antennal lobes (AL) of adult females, we analyzed the neural activity elicited by CO_2_ alone, CO_2_ with a host odor blend, and CO_2_ combined with individual components of the host odor blend (Fig. 3E).

We found a significant modulation of the neural responses to CO_2_ by the presence of host VOCs in the background (Fig. 3F,G). In particular, some neural units that were unresponsive or showed weak responses to CO_2_ in isolation, exhibited heightened excitatory responses when presented with CO_2_ in combination with host odorants such as lactic acid (units 6, 54), nonanal (unit 65), or the blend of host odorants (unit 51) (Fig. 3F). Overall, 19.7% of units significantly increased their firing rate (Type III ANOVA from LMM: F_2,75_ = 5.66, *p* = 0.0051), 21.1% significantly decreased (F_2,80_ = 9.38, *p* < 0.001), and 59.2% did not show any significant change (F_2,225_ = 0.19, *p* = 0.83). Altogether, this resulted in a modulation of the representation of CO_2_ in the neural ensemble by host-related chemicals (Fig. 3G). The addition of lactic acid and nonanal produced the most important shifts while the host odor blend shifted the representation of CO_2_ in a separate direction, suggesting a synergistic effect of chemicals in the blend. Altogether, these results highlight the context-dependent nature of CO_2_ encoding.

Next, we investigated whether CO_2_ modulates plant and host VOCs’ representation in a group of mosquitoes that show strong body size dependency in their behavioral response to these cues. Responses to both host and plant odor blends and their components were compared in two olfactory backgrounds: carbon dioxide and a humidified stream of clean air. The results revealed that CO_2_ did not affect responses uniformly (PERMANOVA, F_1,18_ = 1.09, *R*_2_ = 0.057, *p* = 0.35, based on 999 permutations) and did not differentially modulate host-from plant-associated chemicals (Permutation test, *p* = 0.48), but interacted in an odor-specific manner to modulate how odors are processed in the AL (Fig. 3H,I). This suggests that CO_2_ selectively fine-tunes olfactory processing. Supporting this idea, the representation of the plant blend, distinct from the host blend in the absence of CO_2_, shifted to be significantly closer to the host blend in the presence of CO_2_ (by 5.17 PCA units; permutation test, *p* = 0.046; Fig. 3H). The next odorant showing the largest, albeit non-significant, modulation is benzaldehyde, whose representation tended to be closer to the host blend in the presence of CO_2_ (by 1.69 PCA units; permutation test, *p* = 0.29; Fig. 3I). Altogether, our results show a bidirectional interaction between CO_2_ and other VOCs, highlighting that the relationship between CO_2_ and odorants is more complex than previously understood. CO_2_ is not only an activator of mosquito behavior that places females in a “persistent host-seeking state” (31), but it also modulates host-seeking behavior by modulating the processing and integration of critical host and plant odors. Behavioral assays revealed that virgin females, unlike their mated counterparts, do not respond to CO_2_ alone but exhibit size-dependent responses when exposed to host blend (Fig. S2A1-3,C1-3). Together, these effects likely allow mosquitoes to optimize their likelihood of locating relevant resources in dynamic olfactory and physiological contexts.

### Seven hub-genes regulate body size-dependent variation in the mosquito transcriptome

To decipher the molecular correlates associated with the body-size- and mating-status-dependent effects we observed, we conducted RNAseq analyses. Specifically, we compared the transcriptome of the whole heads of small and large females, either mated or virgin, as the head houses key tissues—including the brain and peripheral sensory organs—involved in olfactory processing, feeding, and reproductive behaviors, allowing us to capture gene expression differences relevant to both body size and mating status (Fig. 4A1-A3).

**Fig. 4.**
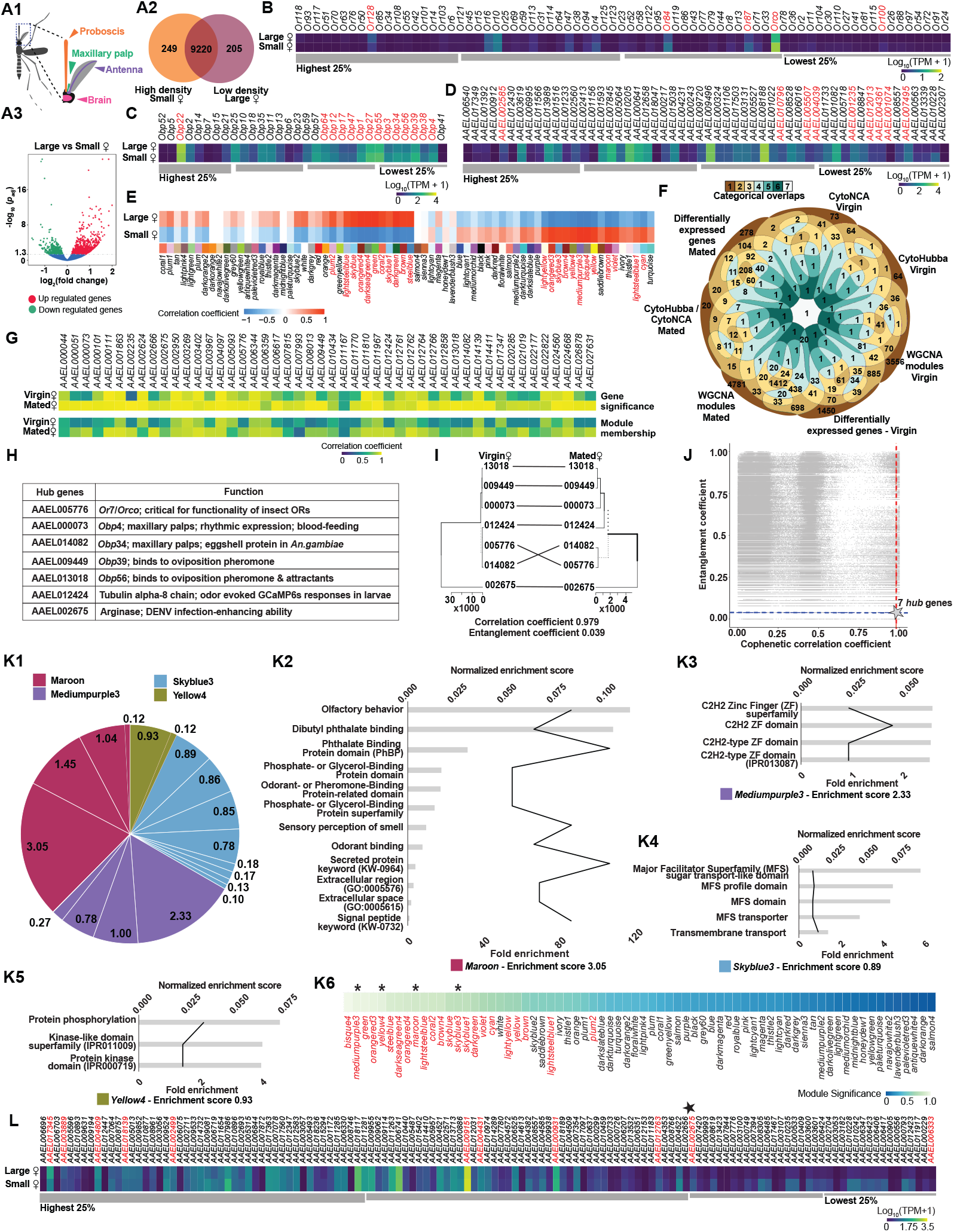
Transcriptomic profiling reveals hub genes driving body size-dependent gene expression in mosquitoes. **(A1)** Schematic of the whole mosquito head subjected to transcriptomic profiling to compare gene expression between large and small female mosquitoes. **(A2)** Venn diagram showing the number of unique and co-expressed genes in large and small mated females. **(A3)** A volcano plot illustrates effect sizes and statistical significance for up- and downregulated genes. **(B-D)** Differential transcript abundance is presented for key gene categories: odorant receptors **(B)**, odorant-binding proteins **(C)**, and immune function-related genes **(D)**. Gene names in red denote transcripts that are significantly differentially expressed between size classes (adjusted *p* ≤ 0.05). **(E)** Heatmap of module-trait correlations from Weighted Gene Co-Expression Network Analysis (WGCNA). Each column represents a gene module, and rows indicate correlations for large and small females. Cells are color-coded to reflect the correlation magnitude, with significant modules labeled in red (Pearson *r, p* < 0.05). **(F)** 7-set Venn diagram showing the overlap between gene lists identified by supervised (differentially expressed genes, DEGs) and unsupervised approaches (WGCNA modules and PPI networks via CytoHubba and CytoNCA). This includes four gene lists for virgin females and three for mated females, highlighting critical body size-dependent and mating status-independent genes (see methods for details). Diagram design adapted from Santiago Ortiz’s 7-set Venn diagram (https://moebio.com/research/sevensets/). **(G)** Heatmap of gene significance (GS) and module membership (MM) for the 48 genes identified in panel F. Each column represents a gene, and rows correspond to virgin and mated females. Cells are color-coded to indicate the magnitude of correlation. **(J)** Scatter plot of cophenetic correlation (X-axis) and entanglement coefficient (Y-axis) for 2.5 million randomly selected 7-gene combinations out of 73.6 million possible combinations (48C7) for the 48 genes listed in panel G. The original seven hub genes, listed in panel H, are visualized in panel I, where their correlation values are denoted by a star. Of the sampled combinations, 0.18% outperformed the hub genes (points below the dashed blue line and to the right of the dashed red line), with 77.3% of these top-performing combinations containing at least one hub gene. **(K)** GO enrichment analysis of gene modules identified in panel E, which contain the 7 hub genes (Maroon, Mediumpurple3, Skyblue3, and Yellow4). Subpanels (K2–K5) summarize GO terms with fold enrichment and normalized enrichment scores. (K6) WGCNA modules ranked by module significance, with modules containing hub genes marked by asterisks. Module names shown in red indicate those with statistically significant correlations to body size, as identified in panel E. **(L)** List of genes encoding the salivary proteome, with genes showing significant differential expression between large and small females highlighted in red. The hub gene, *AAEL002675*, encoding for *arginase* is denoted by a star.

To characterize the body size-mediated effects on the expression of chemosensory-related genes, we compared the expression of genes encoding odorant receptors (ORs), odorant binding proteins (OBPs), and salivary gland proteins (SGPs) (32–35). Out of the 131 ORs characterized in *Ae. aegypti* (36), we identified 73 OR transcripts and five of them were differentially expressed between large and small-sized females (DESeq2 Negative binomial model: *p* < 0.05; Fig. 4B). Among them, *Or128* was significantly upregulated (log_2_ fold change = 1.071, *p*_*adj*_ < 0.05), while *Or84, Or87, Orco*, and *Or100* were significantly downregulated in large females relative to their small counterparts.

We also detected the transcripts encoding 32 of the 66 OBPs characterized in *Ae. aegypti* (37). The transcripts of 14 OBPs showed significant differences (DESeq2 Negative binomial model: *p* < 0.05; *p* < 0.05; Fig. 4C). Among them, only *Obp22*, overexpressed in large females, has been deorphanized, binding long-chain fatty acids (38) and playing roles in host-seeking, male reproduction, and viral susceptibility (39). OBP1 has been structurally characterized with a pH-sensitive binding pocket, suggesting a role in odorant transport, while OBP10 has been functionally linked to host-seeking, reproduction, and arbovirus transmission, with knockouts showing impaired ZIKV and DENV dissemination (40). Besides these chemosensory genes, we also detected 54 transcripts encoding salivary gland proteins, and nine of them were differentially expressed (Fig. 4D).

Given the complexity of body size as a polygenic trait (41–43), we employed both supervised and unsupervised analyses to identify body-size-dependent variations in gene expression. Two unsupervised, network-based methods were particularly instrumental in uncovering these patterns. First, a Weighted Gene Correlation Network Analysis (WGCNA) identified 4,781 genes, out of a total of 14,909, whose expression states covaried with body size (Fig. 4E). Second, a Protein-Protein Interaction (PPI) network analysis highlighted 20 genes among the 278 differentially expressed genes (DEGs) as core members of protein networks influenced by body-size-dependent variations (Fig. S4 M1-M4). Unlike conventional PPI analyses that rely on proteomics data, this approach maps transcript-level information onto known protein interaction networks, assuming that differentially expressed transcripts correspond to functionally relevant proteins (44, 45). Although transcript abundance may not always reflect abundance and although post-translational modifications can alter protein interactions, we used gene expression as a proxy for potential molecular interactions, consistent with approaches used in prior studies (46–50). This allows for a biologically meaningful assessment of gene networks underlying body-size-dependent traits, revealing functionally co-regulated pathways that may drive mosquito physiological adaptations.

Genes uniquely expressed in either virgin or mated females were classified as mating-status-dependent. In contrast, those differentially expressed between large and small females, regardless of mating status, indicated body-size-dependent and mating-status-independent effects. Using a combination of supervised and unsupervised approaches, we employed an integrative framework to analyze overlaps across seven analytical categories: four for virgin females—DEGs, WGCNA outputs, and two distinct PPI network analyses (CytoHubba and CytoNCA)—and three for mated females – comprising DEGs, WGCNA outputs, and a single PPI category given that CytoHubba and CytoNCA produced identical outputs for mated females.

Through this framework, we visualized overlaps across categories using a comprehensive Venn diagram, which highlighted 127 unique combinations of overlaps, systematically evaluating intersections of supervised and unsupervised approaches (Fig. 4F). Combinations involving five or more categories were particularly significant, as they ensured that at least one category from mated females was included. This approach identified seven “hub” genes, *i*.*e*., putative central nodes in gene regulatory networks underlying physiological, sensory, and behavioral differences between small and large females (Fig. 4G-I). These genes were identified through the intersection of high-confidence criteria across multiple methodologies (see methods for details). Specifically, we analyzed correlation coefficients and entanglement coefficients between the dendrograms of transcripts from virgin and mated females (Fig. 4J). The hub genes exhibited high correlation coefficients (i.e., preserved pairwise distances between the data points) and low entanglement coefficients (*i*.*e*., good alignment), demonstrating their stability across mating statuses. Among the 48 candidate genes identified through this multi-threaded approach (Fig. 4G), only 0.18% of 7-gene combinations outperformed the hub genes in robustness metrics, and 77.3% of these outperforming combinations included at least 1 of the hub genes, further reinforcing their significance (Fig. 4H).

The enrichment analysis of the four WGCNA modules containing the seven hub genes (assigned arbitrary color names, by convention) revealed key molecular pathways underlying body size-dependent variations in mosquito olfactory behavior (Fig. 4K1). The *Maroon* module (which includes hub genes *AAEL005776, AAEL014082, AAEL009449*, and *AAEL013018*) is enriched for genes involved in odorant detection and binding, suggesting a role in peripheral olfactory processes such as signal transduction in antennae and maxillary palps (Fig. 4K2) (51–53), such as including odorant-binding proteins (OBPs), which facilitate the detection of cues essential for host-seeking behavior (52, 54–56). The *Mediumpurple3* module (which includes a hub gene *AAEL012424*) shows enrichment for genes encoding C2H2-type zinc finger transcription factors, which are key regulators of gene expression involved in metabolic control (Fig. 4K3) (57, 58). C2H2 zinc finger proteins, including Krüppel-like factors, regulate key physiological functions such as energy homeostasis and stress responses (59, 60). As transcriptional regulators, they likely orchestrate metabolic adaptation to size-specific variations, ensuring coordinated physiological adjustments. This aligns with findings in other taxa, where size-associated traits like longevity, reproduction, and immunity are governed by complex transcriptional networks (61–63). The *Skyblue3* module (which includes the hub gene *AAEL002675*) highlights the role of transmembrane transport processes, with significant enrichment for genes encoding transporters from the Major Facilitator Superfamily (MFS), which mediate nutrient and ion transport (Fig. 4K4) (64) critical for maintaining cellular homeostasis and supporting the higher metabolic demands of larger females (65).

The *Yellow4* module (which includes the hub gene *AAEL000073*) is enriched for genes involved in protein phosphorylation and intracellular signaling, suggesting that these pathways regulate growth, reproduction, and metabolic adaptation (Fig. 4K5) (66–71). Protein kinases, which mediate phosphorylation-based signal transduction, enable mosquitoes to respond dynamically to environmental and physiological changes, providing flexibility to cope with variable conditions (72–74). These signaling cascades are crucial for integrating growth cues, hormonal regulation (*e*.*g*., ecdysone signaling), and reproductive activity (75), key determinants of vector competence.

Together, these findings highlight a robust molecular framework that drives body size-dependent adaptations, shaping key phenotypic traits such as olfactory sensitivity, reproduction, and metabolic regulation. The hub genes within some of the most significant WGCNA modules (identified through high module significance scores) are integral to maintaining regulatory networks that vary with body size, highlighting their importance in coordinating key physiological and behavioral adaptations (Fig. 4K6).

Functionally, one of the seven hub genes remains uncharacterized, offering exciting opportunities to further explore novel mechanisms underlying mosquito behavior and physiology. Five of the remaining six hub genes are implicated in olfaction, a critical sensory modality mediating host-seeking behavior, including *Orco*, an obligatory subunit of ligand-binding olfactory receptors (ORs). Finally, the last hub gene encodes arginase, a salivary protein implicated in dengue virus infection 76, suggesting a potential connection between body size and pathogen transmission. This link is further supported by the differential expression of several genes associated with the salivary gland proteome (Fig. 4L), indicating the importance of body size in both sensory and vectorial functions.

### Transstadial effects of larval competition affect vector transmission potential of mosquitoes

Given the implication of the *arginase* hub gene in arboviral infection (76), we quantified epidemiologically important physiological traits in small, large, and colony-derived females. Specifically, we compared Zika virus (ZIKV) transmission potential across these groups, and the survival probability and timing of oviposition, under sugar- and water-fed conditions, with and without exposure to ZIKV-infected bloodmeal (Fig. 5). In sugar-fed mosquitoes, ZIKV infection significantly reduced survival across all three groups (Hazard ratio (HR) = 2.64, *p* < 0.001; LT_50_: uninfected=28 days, infected=13 days), indicating significant physiological costs of infection (Fig. 5A). Interestingly, small-sized females exhibited overall higher survival compared to colony and large females (HR = 0.28, *p* < 0.001, LT_50_: small = 34.5 days, large = 19.0 days, colony = 8.0 days, Fig. 5A1). The survival of uninfected large females was similar to that of uninfected colony-derived mosquitoes (HR=1.56, *p* > 0.05, Fig. 5A2,A3). However, ZIKV infection significantly reduced survival in colony females (HR = 0.38, *p* < 0.01), who had the shortest lifespan following infection (LT_50_ =8.0 days, Fig. 5A3), whereas infection had little to no effect on large females (HR=0.61, *p* > 0.05, Fig.5A2). In water-fed mosquitoes, ZIKV infection had no significant effect on the survival of large-sized (HR = 1.10, *p* > 0.05, LT_50_ = 8.0 days, Fig.5B1), small-sized (HR = 1.06, *p* > 0.05, LT_50_ = 7.0 days, Fig.5B2), or colony-derived females (HR = 0.56, *p* > 0.05, LT_50_ = 6.0 days, Fig.5B3). Water-fed mosquitoes, with their much shorter lifespan, likely did not live long enough for ZIKV to complete its extrinsic incubation period (typically 3-14 days), limiting the time the virus had to exert a physiological toll. As a result, infection-related mortality may have been masked by the naturally short lifespan of water-fed mosquitoes, as they died before the virus could significantly affect their physiology.

**Fig. 5.**
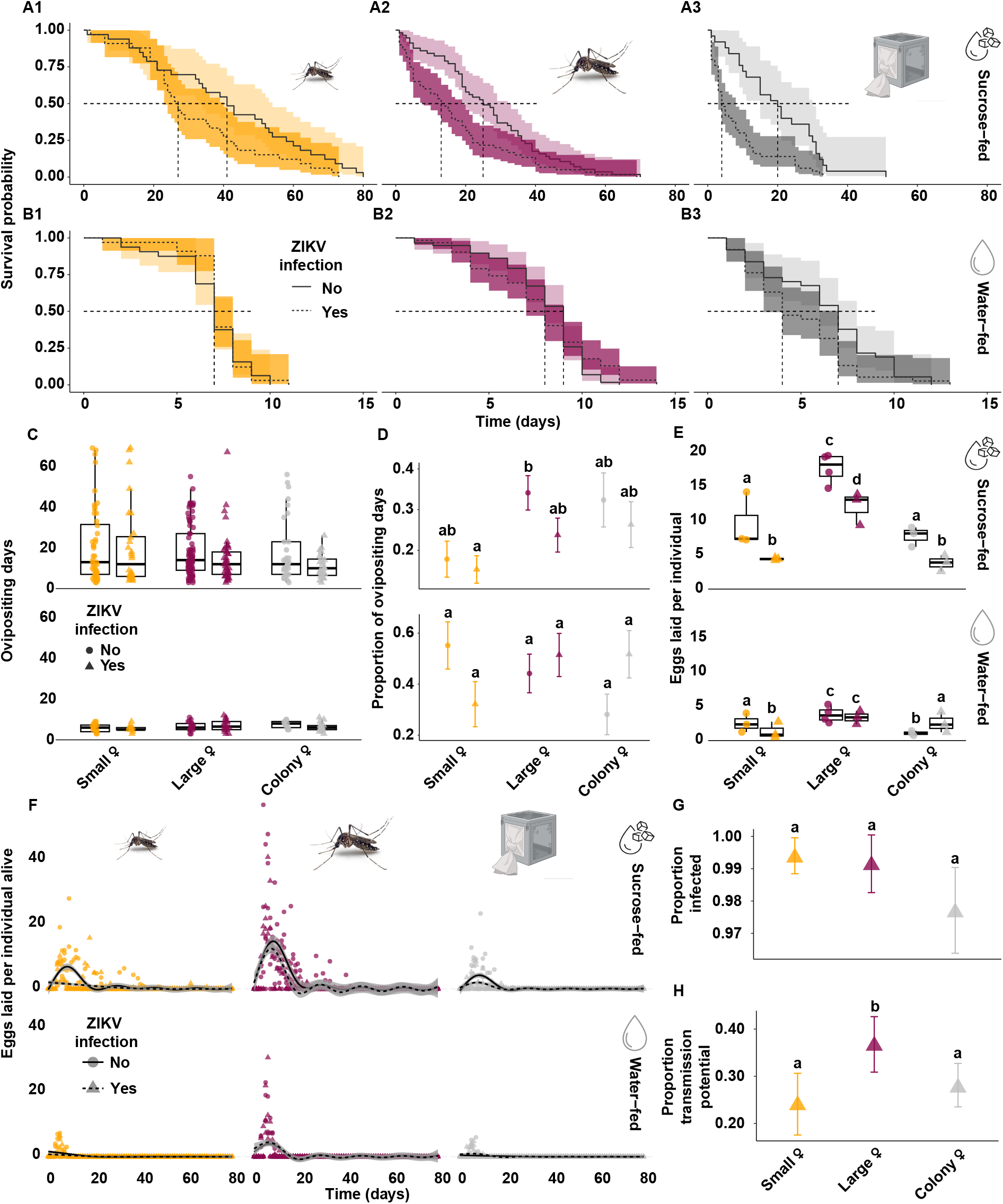
Body size modulates the effects of ZIKV infection and transmission potential. **(A–B)** Survival probability of blood-fed small, large, and colony-derived females with (solid lines) and without (dashed lines) ZIKV infection. ‘No ZIKV infection’ indicates mosquitoes that were fed on a non-infectious bloodmeal and were not exposed to the virus. Longevity was tracked from blood-feeding until death, with females provided ad libitum 10% sucrose **(A1–A3)** or deionized (DI) water **(B1–B3)**. Survival data were analyzed using Cox’s proportional hazards model, with dashed lines indicating 50% survival probability. **(C–D)** Oviposition behavior monitored from blood meal to death, comparing ZIKV-infected (triangles) and non-infected (circles) small, large, and colony-derived females. **(C)** Number of ovipositing days recorded for females fed either 10% sucrose (top panel) or DI water (bottom panel) and analyzed using generalized linear mixed-effects models with Poisson errors. **(D)** Proportion of days post-blood meal in which egg-laying was observed, analyzed using generalized linear mixed-effects models with binomial errors. **(E)** Number of eggs laid per female through their lifetime after one bout of blood feeding, analyzed using a Poisson GLM with a log link and a log-transformed offset for the number of mosquitoes. The model included larval density, infection status, and feeding regime as interacting predictors. **(F)** Number of eggs laid per surviving females daily through their lifetime, assessed using a Gaussian GLM with identity link, incorporating larval density, infection status, and their interaction as predictors. **(G)** Proportion of mosquitoes with ZIKV detected in the midgut ten days after an infectious blood meal. **(H)** Proportion of mosquitoes capable of transmitting the virus, determined by detecting ZIKV in saliva, ten days after the infectious blood meal. Proportions in panels G and H were analyzed using generalized linear models with binomial errors. Error bars in all panels represent mean *±* s.e.m. Exact sample sizes and summary statistics are provided as source data.

ZIKV infection did not affect the number of days females, both sugar- and water-fed, invested in ovipositing (Fig. 5C; Negative binomial GLMM: *χ*^2^ = 0.61, *p* > 0.05). Previous studies indicate that arbovirus infections can affect fecundity, but these effects are often resource-dependent (77). In this study, larval density significantly influenced the effect of ZIKV infection on ovipositing days in water-fed females (Fig. 5D bottom panel; Binomial GLMM: *χ*^2^ = 6.63, *p* < 0.05) but not in sugar-fed females (Fig. 5D top panel; Binomial GLMM: *χ*^2^ = 0.57, *p* > 0.05). However, post-hoc pairwise comparisons showed no significant differences between treatment groups (all *p* > 0.05), suggesting that while larval density modulates the relationship between infection and oviposition, no single group exhibited a distinct effect. However, infected and sugar-fed females laid significantly fewer eggs than their non-infected counterparts (Fig. 5E top panel; Poisson GLM: *χ*^2^ = 14.73, *p* < 0.001). Water-starved females laid significantly fewer eggs than sugar-fed females (Fig. 5E; Poisson GLM: *χ*^2^ = 20.96, *p* < 0.001), and ZIKV infection significantly reduced this number in small females (Fig. 5E bottom panel; *z* = 3.72, *p* < 0.01), had no effect on large females (Fig. 5E bottom panel; *z* = 0.84, *p* > 0.05), and significantly increased this number in colony females (Fig. 5E bottom panel; *z* = -3.20, *p* < 0.05). Oviposition timing differed significantly between feeding treatments (Fig. 5F; Linear mixed-effects model: *χ*^2^ = 43.51, *p* < 0.001), with sugar-fed mosquitoes ovipositing later (17.2 days, 95% CI:13.3–21.1) than water-fed mosquitoes (6.5 days, 95% CI:3.3–9.6). ZIKV infection had no significant effect on oviposition timing (Fig. 5F; Linear mixed-effects model: *χ*^2^ = 3.75, *p* > 0.05), and most individuals laid eggs within 20 days post-blood meal.

Regarding infection and transmission, adult body size did not significantly affect the ability of mosquitoes to become infected with ZIKV (Fig. 5G; Generalized linear model; *p* > 0.05). Infection rates were consistently high and comparable among small, large, and colony-derived females following exposure to the virus (Fig. 5G). However, large females exhibited a significantly higher transmission potential (i.e., the virus was detected in the saliva of 36.4% of them) than colony-derived (27.5%) and small females (24%) (Fig. 5H).

These findings show a body-size dependency in ZIKV transmission, with larger females acting as more efficient vectors compared to smaller or colony-derived females (Generalized linear model; *z* = 2.27, *p* < 0.05). These effects are most likely due to the greater energy reserves of larger mosquitoes and their ability to support more efficient viral replication and dissemination through the salivary glands, as documented in other arbovirus studies (78).

### Transstadial effects shape mosquito demography

Mosquitoes reared under low-density conditions exhibited consistently higher net reproductive rates (*R*_0_) than their high-density counterparts (Fig. SI; F_1,10_ = 21.69, *p* < 0.001), further validating that resource limitation during larval development constrains reproductive output (79–82). Low-density mosquitoes also showed higher intrinsic growth rates (*r*) (range: 0.11-0.16) than the high-density cohort (range: 0.06-0.11) (Fig. S5; F_1,10_ = 23.95, p<0.001). This finding aligns with broader ecological studies that have demonstrated the impact of density on life-history traits in insects (83–86). The strong correlation between estimated (*r’*) and observed growth rates (*r*) validates the life table model, confirming its accuracy in capturing key demographic parameters. Interestingly, cohort generation time (*T*_*c*_), defined as the average interval between the hatching of an individual and the hatching of its offspring, did not differ significantly between density treatments (Fig. S5; F_1,10_ = 1.07, *p* > 0.05), indicating that the underlying physiological mechanisms governing reproductive timing are buffered against environmental variation, whereas the ability to sustain offspring production is far more susceptible to competitive pressures. This is consistent with previous studies that have shown that reproductive timing, rather than total reproductive output, can remain stable when sufficient resources are available (13, 87).

### Transmission modeling elucidates the importance of body size-associated traits and informs intervention strategies

Incorporating mosquito traits, such as variation in body size as the outcome of varied larval conditions, into transmission models provides a more informative framework for assessing risk of disease outbreak (Fig. 6A1-3,C1-3). For equivalent parameter sets, our detailed Larval Mass Model (LMM), shown in Fig. 6A,C, predicted a smaller outbreak, with a slightly delayed peak, compared to the Ross Macdonald Model (RMM), a standard for mosquito-borne disease modeling (88) (Fig. 6B). When the life history of the mosquito was included and informed by the present experiments, competition in the larval stage led to a larger fraction of small mosquitoes, which consequently were less competent for Zika as adults. While the RMM used the average probability of transmission by mosquito size, the actual distribution of mosquito sizes, and its skew towards smaller mosquitoes in the LMM, slowed the outbreak and led to smaller outbreak size. Furthermore, when comparing larval populations experiencing conditions resulting in low- and high-density populations, outbreak size was larger when population density was lower and, thus, composed of larger mosquitoes that allow for more transmission (Fig. 6D).

**Fig. 6.**
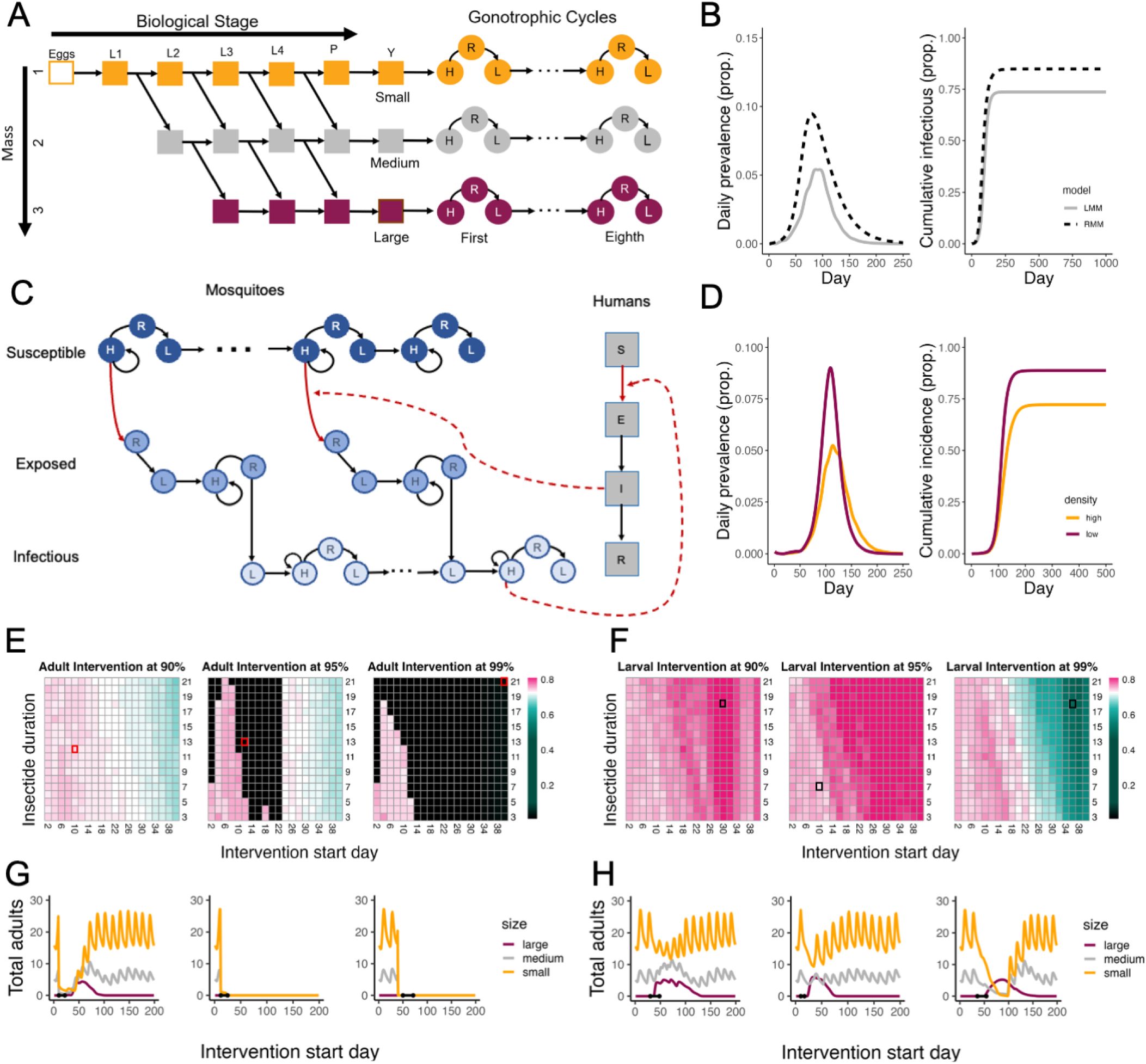
Results from Model Simulations. **(A)** Larval model schematic with population separated by mass. L1–L4, P, Y, H, R, and L represent larvae, pupae, pre-gonotrophic, host-seeking, resting, and egg-laying stages, respectively. **(B)** Comparison of epidemic curve and final proportion of humans infected during outbreak for Ross Macdonald model and detailed larval mass model with transmission. **(C)** Transmission model schematic. H, R, and L represent host-seeking, resting, and egg-laying stages, respectively. The human population is represented by a standard SEIR framework: S (Susceptible), E (Exposed), I (Infectious), and R (Recovered). **(D)** Comparison of low density and high density larval scenario for epidemic curve and final proportion infected in the detailed larval mass model with transmission. **(E**) Comparison of adult intervention at various levels (left to right: 90%, 95%, 99%) starting on various days (x-axis) for various lengths (y-axis). Time series plots for highlighted cells are shown in **(G). (F)** Comparison of larval intervention at various levels (left to right: 90%, 95%, 99%) starting on various days (x-axis) for various lengths (y-axis). Time series plots for highlighted cells are shown in **(H)**. For panels **(E)-(F)**, white represents baseline level of infection, equivalent to the no intervention scenario in gray in (B). The pink colors indicate “more infections than baseline” and turquoise colors is “fewer infections than baseline”. **(G)** Time series plots of adult mosquitoes by mass group showing different start time and durations for adult interventions (left to right: 90% starting on day 10 with a duration of 12 days; 95% starting on day 12 with a duration of 13 days; 99% starting on day 40 with a duration of 21 days). **(H)** Time series plots of adult mosquitoes by mass group showing different start time and durations for larval interventions (left to right: 90% starting on day 30 with a duration of 18 days; 95% starting on day 10 with a duration of 7 days; 99% starting on day 36 with a duration of 18 days).

The type, efficacy, timing, and length of mosquito-targeted interventions during a Zika outbreak alter not only how much but whether the outbreak is attenuated. Overall, adulticides more reliability reduced outbreak size over a wide range of efficacy, timing and length (Fig. 6E), though larvicides if able to be applied with high efficacy had the potential for the greatest reductions in outbreak size (Fig. 6F). However, lower efficacy larvicides almost always increased outbreak size, especially when started later in the outbreak (Fig. 6F). While lower mortality adulticides were effective at reducing the cumulative number of infections during an outbreak (Fig. 6E), with higher efficacy adult interventions, the time at which the intervention started relative to the outbreak impacted intervention effectiveness: earlier intervention may actually have the unintended consequence of higher numbers of infections. This resulted from a decrease in adult population size leading to reductions in larval density and the survival of larger mosquitoes more competent for ZIKV transmission. In contrast, larval interventions later in the outbreak caused the biggest increases as the time lag until reduction in larval density and emergence of large mosquitoes was shorter, coinciding with sharp growth of the outbreak. Overall, the time at which the interventions started was more important for reducing outbreak size than the duration of the intervention, which led to little variation in outcomes for durations of three to twenty-one days.

## Conclusions

Altogether, the present work highlights the profound impact of larval competition on the development and adult traits of *Ae. aegypti*, including survival, reproductive output, host-seeking behavior, olfactory neurophysiology, and vector competence. The stability of generation time across larval densities highlights the resilience of *Ae. aegypti* populations, enabling rapid recovery when larval conditions improve. This echoes the Hydra effect, where partial suppression efforts—such as larval habitat reduction or sublethal insecticide exposure—can inadvertently sustain or even amplify vector populations (89, 90).

Larger females, emerging from low-density larval conditions, show a stronger attraction to host odors and reduced aversion to repellents, while responses to CO_2_ remain consistent across body sizes. Although the tuning of their olfactory sensitivity to host and plant odors is unchanged at the peripheral level, we identified bi-directional interactions between the neural encoding of CO_2_ and resource-specific VOCs, which modulate each other’s representation in the antennal lobes. These central processes fine-tune how these resource-specific cues are processed and integrated, challenging the conventional view of CO_2_ as a mere host-seeking activator (31, 91) and revealing a previously unrecognized complexity in mosquito sensory integration.

Crucially, transcriptomic analyses identified key regulatory hub genes associated with size-dependent traits, offering molecular insights into the transstadial effects through which larval growing conditions shape adult host-seeking behavior and vector competence. These hub genes govern olfaction, metabolism, signal transduction, and salivary gland proteome function, critical processes linking early environmental stress to adult phenotypic performance. Integrating these mechanistic findings into a disease transmission model, we demonstrate that incomplete larval control can paradoxically increase outbreak risk by enabling a subset of survivors to emerge as larger, longer-lived, and more competent vectors.

Classical models that omit transstadial effects—such as the Ross-Macdonald framework—simplify transmission risk by assuming mosquito populations are homogeneous and attribute transmission risk primarily to adult abundance, overlooking size-structured variation in vector competence and host-seeking behavior. In contrast, building off of our previous work (92), we developed a larval-stage model that incorporates mosquitoes’ physiological and behavioral traits linked to body size on disease transmission. This framework allows for the evaluation of intervention strategies targeting the larval or adult stages.

More broadly, our results challenge the long-standing assumption that metamorphosis severs the influence of early-life environments on adult traits in holometabolous insects. Instead, we show that environmental conditions during development leave persistent, gene-regulated signatures on adult phenotypes, with significant ecological and epidemiological consequences. By incorporating these fine-scale biological insights into epidemiological models, we reveal how size-structured heterogeneity in vector competence and host-seeking behavior fundamentally alters outbreak dynamics. As climate change, urbanization, and habitat alteration continue to modify larval environments, understanding the transstadial transmission of environmental effects will be critical for predicting, managing, and mitigating future vector-borne disease risks.

## ACKNOWLEDGEMENTS

We thank Saied “JP” Mirlohi and Shajaesza Diggs for their help in maintaining the *Aedes aegypti* colony. We also thank Daniel Slade, Joshua Benoit, Jake Tu, Igor Sharakhov, and Diane Eilerts for their comments and advice on the analysis of transcriptomic data, Kylie Allen for sharing her vacuum chamber for the synchronization of mosquito hatching, Danny Eanes for technical assistance with custom-made equipment, and The Fralin Life Sciences Institute and the Department of Biochemistry at Virginia Tech for access to shared facilities and equipment. The following reagent was obtained through BEI Resources, NIAID, NIH: *Aedes aegypti*, Strain ROCK, MRA-734, contributed by David W. Severson. This study was partially supported by the National Institute of Allergy and Infectious Diseases of the National Institutes of Health under Award Number R01AI155785 and R21AI166633 (to C.V. for shared incubator space and mosquito activity monitoring), the USDA National Institute of Food and Agriculture, Hatch Research projects VA-160212 (to C.V.), the Center for Emerging, Zoonotic, and Arthropod-borne Pathogens (CeZAP) with pilot funds (to C.V., and L.C.).

## AUTHOR CONTRIBUTIONS

Conceptualization: K.C. and C.V.; physiological and behavioural experiments and analysis: K.C., A.B., and C.V.; electrophysiology and analysis: K.C., S.R., C.L., and C.V.; virology experiments and analysis: K.C., J.M.M., and J.W.L; RNA sequencing and analysis: K.C. and C.V.; epidemiological modeling and visualization: M.W., M.A.R. and L.C.; funding acquisition: K.C., C.V. and L.C.; supervision: C.V., C.L. and L.C.; writing: ; editing: all authors.

## Methods

### A. Mosquito colony maintenance and larval rearing

We sourced *Ae. aegypti* eggs (Rockefeller strain, MR-734, MR4, ATCC©, Manassas, VA, USA) from our laboratory colony maintained at 26°C and 60% relative humidity (RH) under a 12-12h light-dark cycle. Adult mosquitoes were fed *ad libitum* with 10% sucrose and blood fed weekly using an artificial feeder (part # CG-1836-75, ChemGlass, Vineland, NJ, USA) with heparinized blood (Lampire Biological Laboratories, Pipersville, PA, USA) warmed at 37°C using a circulating water bath (RTE-111 Neslab, Thermo Fisher Scientific, Waltham, MA, USA). After blood feeding, mosquitoes were provided with plastic cups (Choice 5.5 oz. Clear Plastic Souffle Cups, Webstaurant, Lititz, PA, USA) half-filled with water and lined with coarse brown seed-germination paper (Anchor Premium Seed Germination Paper, Hoffman Manufacturing Inc., Corvallis, Oregon). Females laid their eggs on the damp paper, which was then removed and dried to prevent hatching. The eggs were stored in a climatic chamber under colony conditions (see above) for at least three weeks before being used in the experiments.

The stored eggs were synchronously hatched in 300 mL of deionized water using a vacuum chamber (set at 21 mm Hg for 60 minutes). Freshly hatched first instar larvae were then distributed in densities of 26 and 78 per 300 mL of nutrient medium stock at 12.5% dilution in the experiment containers (24 fl oz storage containers, McCormick, Baltimore, MD, USA) to simulate ‘low’ and ‘high” intraspecific competition. The stock larval nutrient medium was prepared at 3.3 mg/mL using standard fish food (Hikari Tropic First Bites, Petco, San Diego, CA, USA), incubated at 26C for 24 hours, and used to prepare 12.5% stock dilution. The experiment containers were housed in a climatic chamber set at 26°C and 60% relative humidity (RH) under a 12-12h light-dark cycle.

Mosquito larvae were monitored every 12 hours until pupation. Upon pupation, they were transferred to individually labeled vials containing 10 mL of deionized water and monitored until emergence using a locomotor activity monitor (LAM25, Trikinetics Inc, Waltham, MA, USA). Three infrared beams bisected each vial just above the water level, repeatedly recognized by opposing infrared detectors. Beam breaks were recorded at repeated 60 s intervals to detect emerging mosquitoes (Fig. 1A). We recorded the larval development times and proportion surviving through every stage metamorphosis (four larval instars, pupa, and adult) until emergence. Within 4 hours post-emergence, adult females were weighed using a microbalance (XA105DU, Mettler Toledo, Columbus, Ohio) to quantify their body mass. They were then fed with 10% sucrose solution for five days before being starved for 24 hours on day six until subsequent assays except for quantifying adult longevity and activity. Wet-starvation is known to enhance adult females’ motivation, specifically in host-seeking and blood-feeding contexts (93).

### B. Standardization of larval densities and stock nutrient medium required to simulate heterogenous growing conditions

Larval densities were manipulated to vary *per capita* nutrition availability across treatments, ensuring values comparable to those observed in natural mosquito habitats (16). Prior studies with similar larval densities have demonstrated direct and indirect effects of intraspecific competition on larval and adult traits, including body size (11). To determine the optimal dilution of the stock larval nutrient medium, we recorded the number of mosquito larvae surviving to pupation under low (26 larvae per 300 mL) and high (78 larvae per 300 mL) competition conditions across 10%, 12.5%, and 15% stock medium dilutions. Among these, the 12.5% dilution was selected for the experiments, as it produced sufficient variation in larval development time and body mass at emergence—key proxies for adult mosquito body size (94). This dilution yielded an adequate number of adult females per cohort to support downstream behavioral, electrophysiological, and molecular assays. Additionally, the extent of variation observed in the 12.5% dilution, including differences in larval development and adult female body mass at emergence, was comparable to the variability in larval densities reported in our previous studies and consistent with data from other laboratory investigations on *Ae. aegypti* (11). This reinforces its ecological relevance and experimental robustness of the selected rearing conditions. This standardization ensured that observed differences in adult traits, including host-seeking behavior, olfactory sensitivity, and gene expression, emerged from biologically relevant developmental gradients and were not confounded by nutritional extremes or insufficient replication. As such, the experimental design provides a reproducible and ecologically grounded framework for probing the physiological and behavioral consequences of body size variation in mosquitoes.

### C. Quantification of life history and demographic traits

#### C.1. dult activity and longevity

The activity and longevity of adult *Ae. aegypti* females reared under low and high competition conditions were quantified using a locomotor activity monitor (LAM25, Trikinetics, Waltham, MA, USA, Fig. 1A). Within 4 hours post-emergence, following body mass quantification, individual females were transferred to horizontal glass vials (25 mm × 95 mm) within the activity monitor. Each vial was fitted with a cotton plug saturated with a 10% sucrose solution, replenished as needed, to ensure *ad libitum* access to food. The activity monitor was housed within a climate-controlled incubator (26°C, 60% RH, 12:12 h light-dark cycle) to minimize external disturbances. The LAM25 software continuously recorded infrared beam breaks at 1 min intervals to track locomotor activity throughout the lifespan of each mosquito. Longevity was determined based on the last movement event recorded by the software.

Periods of inactivity were recorded as zeros, resulting in a zero-inflated dataset. To account for this, we calculated the William’s Mean (Mw), a modified geometric mean designed to accommodate zero inflation in activity data for all monitored mosquitoes (95). A pronounced startle response was observed immediately following the transition to darkness, consistent with previous studies (27, 96, 97). Data recorded at the moment of lights-off were thus excluded and interpolated using the arithmetic mean of activity values recorded immediately before and after the transition (27). For longevity analysis, given the known positive correlation between female body size and lifespan, we used Cox’s proportional hazards model with wing length as a covariate. This approach allowed us to estimate the hazard ratio for mortality and assess differences in survival between treatment groups (11, 98).

At the end of the assay, dead mosquitoes were removed, and their wings were dissected for body size measurements. Wings were detached using precision forceps, imaged alongside an ocular micrometer, and measured from the anal lobe to the wing tip to the nearest 0.1 mm using ImageJ software (99).

#### C.2. Blood feeding and oviposition behavior

To quantify blood feeding and oviposition behavior, freshly emerged adult females from three low and one high intraspecific larval competition treatment cohorts were housed in separate mesh cages (BugDorm-1 Insect Rearing Cage, MegaView Science Co., Ltd., Taichung, Taiwan) along with adult males (at a ratio of 3 males per female) reared under standard laboratory conditions.

To enable individual tracking throughout the blood feeding and oviposition assays, each female mosquito was marked with a unique color code using odor-free, waterproof paint (Basics Acrylics Color, Liquitex, Cincinnati, OH, USA) applied to the ventral side of the abdomen. Standardization assays confirmed that paint application had no significant effect on female activity or reproductive behavior.

Between days 6 and 11 post-emergence, adult females in each cage were offered a blood meal daily at ZT 10-12 using the same procedure as for colony rearing. Each blood-feeding session lasted 45 minutes over six consecutive days. Following each session, all engorged females were immediately isolated and identified based on their color-coded markings before being isolated in plastic vials (Flystuff Narrow Drosophila Vials, Genesee Scientific, El Cajon, CA, USA) lined with a moist coarse brown seed-germination paper disc at the bottom, to facilitate oviposition. The age at which they successfully blood-fed was recorded.

The paper lining the vials was moistened daily with 500 *µ*L of deionized water, and oviposition was monitored daily so the first day of egg laying could be recorded for each female. Females were allowed to oviposit for up to 30 consecutive days, after which their wings were dissected and measured. Standardization assays revealed that females deposited the majority of their eggs between 3 and 6 days post-blood feeding. To confirm whether eggs were retained beyond this period, dissections on a subset of females were conducted on day 7 post-blood feeding, revealing that most ovarian follicles were in Christophers’ stages II, III, and IV, with only a few mature eggs persisting in Christophers’ stage V (100). Egg strips from individual females were labeled, dried, and stored in a climatic chamber under the same conditions as the main experiment for 30 days. The number of eggs per strip was manually counted under a microscope (AmScope SM-2 Series, Amscope, Irvine, CA, USA). The egg strips were then immersed in 300 mL of deionized water, and the number of larval progeny hatching over a 14-day period was recorded. Throughout the blood feeding and oviposition assays, adult female mosquitoes had *ad libitum* access to 10% sucrose solution to ensure baseline nutritional support.

The effects of larval competition were assessed by comparing the proportions of blood-fed and ovipositing females across treatment groups using a generalized linear model (GLM) with binomially distributed errors, treating intraspecific competition treatment (‘low’ *vs*. ‘high’) as the independent variable. For each female, we quantified the oviposition latency, defined as the time interval between blood feeding and first egg oviposition, and analyzed it using a linear model (LM) with normally distributed errors. Fecundity was measured as the number of eggs laid in the first gonotrophic cycle, with wing length (a proxy for body size) included as a covariate. The fecundity data were analyzed using a GLM with Poisson-distributed errors to account for count-based variability.

#### C.3. Life-Table construction and population growth estimates

To evaluate the demographic consequences of larval competition, we constructed cohort life tables and estimated key population growth parameters following established approaches on mosquitoes, specifically *Ae. aegypti* (80, 101). These included the net reproductive rate (*R*_0_), cohort generation time (*T*_*c*_), and the intrinsic rate of increase (*r*), calculated by solving the Euler–Lotka equation iteratively using solver add-in in MS Excel (Microsoft 365 v16.93.1). We used ANOVA to assess the effects of intraspecific larval competition on life-table traits. All statistical assumptions were verified and met. All analyses were conducted using R version 4.4.2 (102).

#### C.4 Physiological trait-space analysis

To evaluate how variation in adult body size governs coordinated expression of survival and reproductive traits, we constructed a multivariate, principal component analysis (PCA) that integrates individual-level data on longevity, fecundity, and offspring viability. While these traits reflect discrete life history outputs, their integration offers a composite view of physiological performance. Wing length, log-transformed to serve as a continuous proxy for body size, was used as the primary predictor across all datasets. Generalized additive models (GAMs) were fit to population-level data to characterize nonlinear relationships between body size and each trait: total egg production, oviposition probability, progeny number, proportion of viable offspring, and adult longevity under water- and sucrose-fed conditions. Each model was smoothed using thin-plate regression splines and fitted using restricted maximum likelihood (REML) to balance model flexibility with generalization. Because all traits were not quantified on the same individuals, predictions were generated from the fitted GAMs to estimate missing trait measurements. Linear interpolation was applied within the modeled range, while boundary extrapolation was used for individuals falling outside the empirical range. This procedure enabled estimation of fecundity traits in longevity datasets and vice versa. All estimated and observed trait values were retained for downstream analyses.

The resulting composite dataset was standardized and subjected to principal component analysis (PCA), using egg count, oviposition probability, progeny output, offspring viability, and longevity under both dietary conditions (*i*.*e*., sugar or water fed). Individuals were positioned in a continuous physiological trait space based on the first two principal components.

### D. Behavioral assays quantifying mosquitoes’ host and plant-seeking preferences

The olfactory preferences of *Ae. aegypti* females were assessed using a Y-maze olfactometer as in (27, 103–105). Olfactory stimuli were delivered via polytetrafluoroethylene (PTFE) tubing (McMaster-Carr, Elmhurst, IL, USA) connected to the distal ends of the choice arms. Charcoal-filtered air, flowing at 3 cm.s^*−*1^, was directed through scintillation vials containing either the test odorant or a control solution (mineral oil). To eliminate positional bias, the test stimulus and control were randomly alternated between the choice arms across trials. Experiments were conducted under controlled temperature and humidity conditions optimal for *Ae. aegypti* activity [26 ± 0.5°C, 80% ± 5%RH]. Individual mosquitoes were introduced into the entrance arm and given 5 minutes to make a choice between the two test arms. Their movement and arm selection were recorded to quantify odor preferences. A neutral control test, in which mosquitoes were presented with two clean air streams, confirmed the absence of inherent side bias in the apparatus.

To resolve how host-seeking behavior varies continuously with mosquito body size, we developed a moving window analysis that avoids arbitrary binning and captures high-resolution behavioral variation across the full range of observed phenotypes. Rather than grouping individuals into discrete size classes, which risks losing information or introducing bias, we treated body size as a continuous variable and applied a sliding sampling window across the ordered distribution of individual body masses. Mosquitoes were first ranked in ascending order of body mass. Based on *a priori* power analysis, we determined that a minimum of 43 individuals was required to robustly estimate olfactory preference in the Y-maze assay (observed proportion *p*_1_ = 0.75, reference value *p*_0_ = 0.5, Type I error rate *α* = 0.05, desired statistical power = 0.97). Beginning with the first 43 mosquitoes in the sorted dataset, we advanced the window by one individual at a time, recalculating behavioral metrics at each step, until the final subset encompassed the last 43 individuals in the size distribution. This yielded a series of overlapping subsets, each representing a local window of body sizes.

For each window, we calculated the mean body mass and the olfactory preference index (PI). The PI summarizes the mosquitoes’ relative choice between two odor stimuli, defined as: *PI* = (*n*_*A*_ *− n*_*B*_)*/*(*n*_*A*_ + *n*_*B*_) where *n*_*A*_ and *n*_*B*_ represent the number of mosquitoes choosing odor A and odor B, respectively. The PI ranges from +1 (complete preference for odor A) to -1 (complete preference for odor B), with 0 indicating equal preference between A and B.

By pairing each preference index with the corresponding mean body mass of its window, the moving window analysis generated a continuous behavioral profile across the body size distribution. This approach enabled fine-scale resolution of host-seeking preferences and identification of size-dependent shifts in behavior without relying on predefined size classes.

#### D.1 Formulation of synthetic host odor and plant odor blends

Synthetic odor blends were formulated to replicate the volatile profiles of two ecologically relevant sources: human body odor (host blend) and a representative plant odor (plant blend). Each mixture aimed to match the vapor-phase composition of naturally emitted odors by adjusting the liquid-phase mole fractions of their component compounds, using Raoult’s law. For the host blend, target vapor-phase proportions were derived from previously reported measurements of volatiles in human odor profiles: Nonanal (2.4%), Butyric Acid (1.0%), Lactic Acid (0.1%), and Octenol (1.0%). For the plant blend, target abundances of Octanal, Benzaldehyde, Linalool, and β-Pinene were obtained from key floral volatiles emitted by orchids pollinated by mosquitoes (105–107).

Raoult’s law was used to estimate the partial vapor pressure (*P*_*i*_) of each compound as a product of its mole fraction in the liquid phase (*X*_*i*_) and its pure component vapor pressure 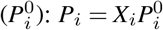

The total vapor pressure (*P*_*total*_) of the mixture was computed as the sum of the individual partial pressures (*P* _*total*_ = ∑ *P*_*i*_), and the vapor-phase composition of each compound was calculated as: *P*_*i*_ / *P*_*total*_

To determine the mole fractions (*X*_*i*_) required to match each blend’s target vapor composition, we implemented a nonlinear optimization routine in Excel Solver. The algorithm minimized the sum of squared deviations between the calculated and target vapor-phase compositions, subject to the constraint that the sum of all *X*_*i*_ equaled 1. This process yielded a liquid-phase formulation for each blend in which the Raoult’s law-derived vapor output closely matched the intended profile.

The host and plant odor blends were prepared using the following chemicals: nonanal (CAS 124-19-6), butyric acid (CAS 107-92-6), L-(+)-lactic acid (CAS 79-33-4), octenol (CAS 3391-86-4), octanal (CAS 124-13-0), benzaldehyde (CAS 100-52-7), linalool (CAS 78-70-6), and *β*-pinene (CAS 127-91-3). All compounds were sourced from Sigma Aldrich, St Louis, MO, USA. For each odor blend, 50X stock solution were prepared following the compositions detailed in Table S5 (host odor blend) and Table S6 (plant odor blend). These formulations were designed to achieve specific mole fractions in the liquid phase, approximating the natural ratios found in human and floral scent emissions, respectively. From the host odor composition, the calculated partial vapor pressures were 0.157 mm Hg (Nonanal), 0.064 mm Hg (Butyric Acid), 0.033 mm Hg (Lactic Acid), and 0.072 mm Hg (Octenol), yielding a total vapor pressure of 0.3259 mm Hg for this solver-adjusted system. The resulting vapor-phase relative abundance was 0.482 (Nonanal), 0.198 (Butyric Acid), 0.100 (Lactic Acid), and 0.220 (Octenol), closely matching the target values from human scent. For the plant odor blend, partial vapor pressures were 0.117 mm Hg (Octanal), 0.100 mm Hg (Benzaldehyde), 0.117 mm Hg (Linalool), and 0.255 mm Hg (β-Pinene), resulting in a total vapor pressure of approximately 0.589 mm Hg. The corresponding vapor-phase relative abundance was 0.198 (Octanal), 0.170 (Benzaldehyde), 0.198 (Linalool), and 0.433 (β-Pinene), matching floral emission profiles (105–107).

#### D.2 Behavioral trait-space analysis

To evaluate how body size variation influences olfactory behavior across experiments involving different odorant choices (Fig. 2), we constructed a multidimensional behavioral trait space using the preference index data from all the olfactometer assays involving colony-bred females. In those olfactometer assays, each female was tested in only one of seven binary choice assays. To capture cross-context behavioral variation at the individual level and enable multivariate comparisons, we used body mass at emergence, measured for all mosquitoes, as a shared predictor to estimate how each individual would respond across all experimental contexts.

For each odor context, we fit a generalized additive model (GAM) to the population-level response data, modeling preference index as a smooth function of log-transformed body mass. Each fitted model was then used to predict responses for every individual across all seven odor contexts. The result was a complete individual-by-context matrix comprising one observed and six predicted values per mosquito. This matrix was standardized and subjected to principal component analysis (PCA) to identify dominant axes of behavioral variation. The first two principal components were used for k-means clustering (k=3) to classify individuals into discrete behavioral phenotypes. Cluster separation was assessed using ANOSIM, and body mass distributions were compared across clusters to evaluate whether olfactory strategies were structured by size variation.

### E. Electrophysiology assays

#### E.1. Mosquito preparation for Electroantennogram (EAG) recordings

Electroantennograms recordings were conducted following the procedure outlined by Lahondère (107). Prior to the experiment, 6–10 days post-emergence mosquitoes were starved for up to 12 hours. Individuals were then cold-anesthetized and the distal segments of both antennae were trimmed and the decapitated mosquito head was mounted onto the reference electrode. The recording electrode was then positioned to enclose the antennae tips. Finally, the preparation was placed within a Faraday cage to minimize electrical noise during data acquisition. Odorant stimuli were prepared in advance by diluting test compounds to desired concentrations (commonly 0.1% or 1%) in mineral oil as the solvent. Glass Pasteur pipettes containing 10 *µ*L of stimuli solution were sealed and allowed to equilibrate for approximately 10 minutes before use. During recordings, a continuous flow of purified, humidified air (140 mL/min) was directed over the antennae to maintain a stable environment. Odorant stimuli were introduced by a 1 second long pulse of the odor-laden air (15 mL/min) into the main airstream. Stimuli were delivered in a randomized order. A vacuum line positioned approximately 20 cm from the preparation ensured rapid removal of odorants following each pulse, preventing adaptation or lingering effects. Control stimuli, consisting of the solvent alone, were interspersed between test odorants to control for mechanical responses. Additionally, a positive control of 10% benzaldehyde, was presented at the end to confirm the responsiveness of the preparation.

Electrophysiological signals were amplified (*x*100), filtered (0.1–500 Hz), and digitized at 50 Hz using WinEDR (University of Strathclyde, Glasgow, UK). Baseline signal stability (< 0.01 mV amplitude fluctuation) was verified prior to each recording. Odor-evoked responses were baseline-corrected by subtracting the corresponding mineral oil control response. Responses to mineral oil (negative control) and benzaldehyde (positive control, 10^*−*1^ concentration), normalized to the mineral oil baseline, were compared across larval density groups using a one-way ANOVA. Statistical comparisons across odorants and treatment groups were performed using two-factor ANOVA with Tukey’s post-hoc test and Bonferroni correction (*p* < 0.05).

#### E.2. Mosquito preparation for extracellular recordings

To quantify the olfactory encoding in the antennal lobe of *Ae. aegypti* mosquitoes, we performed extracellular recordings on adult females aged 6–10 days post-emergence as in (104). Although extracellular recordings are limiting our ability to directly compare small and large mosquitoes, recording from females with body masses ranging around the inflection point for significant host-seeking preference in the olfactometer assay allowed us to capture relevant neuromodulatory effects (Fig. 3H). The responses of 75 neural units across 13 individual mosquitoes were collected. Prior to experimentation, mosquitoes were immobilized by brief exposure to cold anesthesia on ice. Each mosquito was then positioned on a custom-built 3D-printed holder, securing the head while allowing free movement of the antennae (Fig. 3D). Under a stereomicroscope (AmScope SM-2 Series), a small incision was made in the cuticle at the base of the antenna, below the Johnston organ to expose the antennal lobe. Quartz glass capillaries (1.0 mm outer diameter, 0.7 mm inner diameter, Sutter Instrument, Novato, California), pulled to 5 *µ*m diameter tips using a Sutter P-2000 laser puller and filled with 0.15 M sodium chloride solution, served as a recording electrode inserted into the antennal lobe. A reference electrode was placed in the thorax to complete the circuit.

Odorant stimuli were prepared following the methods described for EAGs in the previous section. Neural signals were amplified (*x*10,000) and band-pass filtered (0.3–3 kHz) on an A-M Systems Model 1800 amplifier (A-M Systems Inc, Sequim, WA, USA). Signals were digitized at a sampling rate of 20 kHz and recorded using WinEDR (University of Strathclyde, Glasgow, UK). Spike sorting was performed in Plexon Offline Sorter (Plexon Inc, Dallas, TX, USA) and perievent analysis in NeuroExplorer (NEX Technologies, Colorado Springs, CO, USA). Two pulses of each odorant stimuli were delivered to each preparation. Response magnitudes across odorant stimuli were compared in *R*, by means of principal component analysis. To test for differences in firing rates across conditions (e.g., before, during, after host blend introduction), we fit a linear mixed-effects model (LMM) with firing rate as the response variable, condition (pre-, during, post-) as a fixed effect, and random intercepts for neuron identity to account for repeated measures. Significance of fixed effects was assessed using Type III ANOVA with Satterthwaite’s method for approximating degrees of freedom.

### F. Transcriptomic analysis

#### F.1. Tissue dissection and RNA extraction

To investigate differential gene expression, heads of adult female *Ae. aegypti* from low- and high-density larval treatments were collected on day 6 post-emergence, following a 24-hour starvation period. To ensure circadian consistency, dissections were performed between zeitgeber time (ZT) 10–12. Cold-anesthetized mosquitoes were placed under a dissecting microscope, and their heads were excised using sterile dissection scalpels. The dissected heads were immediately flash-frozen in liquid nitrogen and stored at -80°C until further processing.

For RNA extraction, four biological replicates per treatment group were processed, with each replicate consisting of approximately 40 pooled heads. This pooling strategy was designed to minimize inter-individual variability while maintaining biological relevance. To ensure comparability across replicates, head samples were evenly distributed to account for body size variation within each treatment cohort. Total RNA was extracted using the RNeasy Mini Kit (Qiagen, USA) following the manufacturer’s protocol, which included on-column DNase I treatment to remove genomic DNA contamination. The integrity of extracted RNA was assessed using a NanoDrop 2000 spectrophotometer (Thermo Fisher, USA) by measuring A260/A280 and A260/A230 ratios, ensuring high purity. RNA samples with a RIN (RNA Integrity Number) score greater than 7.0 were considered suitable for downstream sequencing. Extracted RNA was stored at -80°C before shipment for sequencing.

#### F.2. Sequencing, read mapping, and gene annotations

RNA-Seq library preparation and sequencing was conducted by Novogene Co., Ltd. (Beijing, China). Before library construction, RNA quality and purity were verified using the Agilent Bioanalyzer 2100 system (Agilent Technologies, CA, USA). RNA degradation and contamination were assessed via 1% agarose gel electrophoresis, while quantification was conducted using the NanoPhotometer® spectrophotometer. For each sample, 1 *µ*g of total RNA was used as input for library preparation using the NEBNext® Ultra™ RNA Library Prep Kit for Illumina® (New England BioLabs, USA), following the manufacturer’s guidelines.

mRNA was isolated using poly-T oligo-attached magnetic beads and fragmentation using divalent cations under elevated temperature. First-strand cDNA synthesis was performed using random hexamer primers and M-MLV Reverse Transcriptase (RNase H-), while second-strand synthesis was carried out using DNA Polymerase I and RNase H. Double-stranded cDNA was then purified, end-repaired, adenylated at the 3’ ends, and ligated to NEBNext Adaptor sequences with a hairpin loop structure for hybridization. Size selection of fragments (150-200 bp) was performed using the AMPure XP system (Beckman Coulter, USA). PCR amplification was conducted with Phusion High-Fidelity DNA Polymerase, universal primers, and index (X) primers. The final libraries were purified and quality-checked using the Agilent Bioanalyzer 2100 system.

Clustering of index-coded samples was performed using a cBot Cluster Generation System, and sequencing was conducted on an Illumina platform using 150 bp paired-end reads. After sequencing, raw reads were filtered to remove adapter sequences, reads containing more than 10% unknown bases, and low-quality reads with an average Q-score below 20. Clean reads were mapped to the *Ae. aegypti* reference genome (AaegL5.0) from the NCBI database using HISAT2 v2.2.1.0, a splice-aware aligner optimized for RNA-Seq data. The featureCounts v1.5.0-p3 software was used to count read numbers mapped to each gene. Gene expression levels were quantified as fragments per kilobase of transcript per million mapped reads (FPKM) to normalize for transcript length and sequencing depth.

#### F.3. RNA-Seq analysis and differential expression analysis

To ensure consistency and reduce technical bias, quantile normalization of read counts was performed before differential gene expression analysis. Genes with expression levels below 1 FPKM were filtered out as they were considered unreliable for statistical analysis. Differential expression analysis between low- and high-density mosquito groups was conducted using DESeq2 v1.14.1, an R-based package designed for count-based RNA-Seq data (108). DESeq2 employs a generalized linear model based on the negative binomial distribution, allowing for accurate variance estimation in differential expression analysis.

The statistical significance of differentially expressed genes (DEGs) was assessed using the Benjamini-Hochberg false discovery rate (FDR) correction. Genes with an adjusted *p*-value (*p*_*adj*_) of < 0.05 and an absolute log_2_fold change (|log_2_FC|) ≥ 1 were considered significantly differentially expressed. The distribution of differentially expressed genes was visualized using volcano plots, and their expression patterns were further examined through hierarchical clustering and principal component analysis (PCA).

To assess biological relevance, functional enrichment analysis of DEGs was conducted using the clusterProfiler package in R. Gene Ontology (GO) term enrichment was performed to identify significantly overrepresented biological processes, molecular functions, and cellular components. KEGG pathway analysis was conducted to determine metabolic and signaling pathways enriched among differentially expressed genes.

#### F.4. Weighted Gene Co-expression Network Analysis (WGCNA)

To explore transcriptome-wide gene co-expression patterns and their relationship with mosquito body size, we employed Weighted Gene Co-Expression Network Analysis (WGCNA). This unsupervised network-based approach clusters genes into modules based on the similarity of their expression profiles. By constructing a gene co-expression network, WGCNA enables the identification of biologically meaningful gene networks and assesses their correlation with phenotypic traits, providing a systematic framework for uncovering body-size-dependent gene expression patterns (109). We followed established WGCNA protocols for network construction, module detection, and trait association, using soft-thresholding to approximate scale-free topology and dynamic tree cutting to identify modules of co-expressed genes (110–113). Module–trait correlations and gene-level associations were evaluated using eigengene-based approaches, with multiple testing correction applied using the Benjamini-Hochberg method (114).

#### F.5. Transcriptome-based Protein-Protein Interaction (PPI) analysis

To investigate the molecular interactions underlying body-size-dependent gene expression patterns, a Protein-Protein Interaction (PPI) network was constructed using the transcriptomic data described above. While PPIs are traditionally derived from protein-level interactions, transcriptome-derived PPIs provide a functional approximation of molecular interactions by mapping differentially expressed genes (DEGs) to their known or predicted protein counterparts (115–118). Given that the number of transcripts does not always directly correlate with protein abundance, this analysis prioritizes the transcriptional landscape, capturing potential regulatory interactions inferred from gene expression data (119).

Despite this distinction, transcriptome-derived PPIs remain a powerful tool for functional annotation, as transcriptional co-regulation often reflects shared biological pathways and upstream regulatory mechanisms that precede protein interactions (48). While transcriptome-based PPI networks do not capture direct physical interactions, they offer a biologically meaningful framework for understanding gene networks that drive mosquito size-dependent adaptations.

The PPI network was generated using the Search Tool for the Retrieval of Interacting Genes/Proteins (STRING v11.0, https://string-db.org). The input consisted of DEGs identified from transcriptomic comparisons of large and small mosquitoes, mapped to their corresponding protein products. STRING integrates known and predicted PPIs derived from experimental datasets, computational predictions, and curated literature sources. Active interaction sources included experimental data, curated databases, co-expression analyses, text mining, and homology-based predictions, with the species limited to *Ae. aegypti*. A combined interaction score threshold of 0.4 was applied to ensure high-confidence interactions, minimizing the inclusion of spurious associations.

The STRING-generated network was imported into Cytoscape (v3.9.1) for further analysis and visualization. CytoHubba and CytoNCA, two widely used Cytoscape plugins, were employed to identify critical regulatory genes within the network. To assess the structural and functional importance of genes within the network, CytoHubba was used to rank genes based on their direct connectivity and influence in highly interconnected regions. Degree centrality was used to quantify the number of direct interactions a gene exhibited, while betweenness centrality was applied to evaluate genes that act as intermediaries between distinct network regions. Maximal Clique Centrality (MCC) was also used to identify genes embedded within densely interconnected network modules. Genes scoring highly in multiple CytoHubba ranking metrics were considered essential regulators of transcriptional responses linked to body-size-dependent traits.

To complement CytoHubba’s emphasis on direct connectivity, CytoNCA was used to conduct a global network topology analysis that captured both direct and indirect gene influence. Closeness centrality was used to measure the efficiency with which a gene could communicate with others in the network. In contrast, eigenvector centrality accounted for both direct connections and the significance of neighboring genes. Betweenness centrality was reassessed to refine the identification of genes that serve as transcriptional mediators, and subgraph centrality was included to determine the extent to which each gene participated in distinct regulatory modules. Unlike proteomics-driven PPI networks, which can be influenced by post-translational modifications and degradation rates, transcriptome-based networks provide insight into transcriptionally co-regulated pathways and gene clusters.

Following individual analyses conducted by CytoHubba and CytoNCA, a filtering step was applied to integrate and refine the selection of critical genes. The top-ranked genes from each approach were compared, and those identified by both methods were prioritized for further analysis. Additionally, genes that appeared exclusively in CytoHubba or CytoNCA were retained if they exhibited strong network influence based on at least one ranking method. This comparative approach balanced the identification of genes with high direct connectivity and those that exert broader influence on network communication.

To identify functionally significant transcriptome-based interaction clusters, highly connected subnetworks were detected using ClusterONE (Clustering with Overlapping Neighborhood Expansion, v1.0). This clustering method allowed the detection of distinct regulatory modules by identifying groups of genes that interact more densely with each other than with the rest of the network. A minimum cluster size of five genes and a minimum density threshold of 0.05 were applied to ensure that the identified clusters represented functionally relevant transcriptional regulatory units. Edge weights, derived from the combined STRING interaction score, were incorporated to refine cluster boundaries. This analysis provided additional insight into potential functional interactions driving body-size-associated transcriptomic variation.

The final network was visualized in Cytoscape, with node sizes scaled according to connectivity, where larger nodes represented genes with a higher degree of interactions. Edge thickness was adjusted based on interaction confidence scores to visually represent inferred transcriptional regulatory strength. Genes identified through CytoHubba and CytoNCA were highlighted based on their ranking to distinguish those that exhibited high influence across multiple centrality measures.

### G. Vector competence, longevity, and extrinsic incubation period (EIP)

All infectious blood feeds of mosquitoes were performed as previously described (120). Briefly, Zika virus (ZIKV; strain PRVABC59) was used to infect Vero cells (Cercopithecus aethiops kidney epithelial cells, ATCC^®^ CCL-81™) infected at a multiplicity of infection of 0.01 four days before the blood meal. Mosquitoes were starved of glucose one day before blood feeding. On the day of the blood feeding, the virus was harvested and clarified by centrifugation. The final blood meal composition consisted of 45% clarified viral supernatant, 5 nM ATP, and the remaining volume was made up of defibrinated bovine blood (7230801, Lampire Biological Laboratories, Pipersville, PA, USA). Backtiters demonstrated that the average infectious dose to the mosquitoes was 2.6 × 10^6^ plaque-forming units (PFU)/mL. Mosquitoes were fed for one hour using the Hemotek membrane feeding system (SP4W1-3, Hemoteck Ltd., Blackburn, UK). After one hour, mosquitoes were temporarily anesthetized by cold anesthesia, and mosquitoes fed to repletion were selected.

For the vector competence assays, mosquitoes were maintained for 10 days with 10% sucrose ad libitum at 28°C, 12:12h (light:dark) photocycle, and 75% relative humidity. On the tenth day post blood feed, mosquitoes were again anesthetized using cold anesthesia, and bodies and saliva were collected. For saliva samples, mosquitoes were mounted on a 20 *µ*L with 10 *µ*L of Type A immersion oil. Force salivation occurred for 30 minutes, after which the oil was dispensed into 50 *µ*L of mosquito diluent (RPMI-1640 media with 10 mM HEPES + 2% FetalPure FBS + 50 *µ*g/mL Gentamicin + 2.5 *µ*g/mL Amphotericin B). For body samples, after forced salivation, mosquitoes were submerged in 250 *µ*L of mosquito diluent with a small metal bead and homogenized at 30 freq/s for 2 minutes using the Qiagen TissueLyser II after freezing (121). To assess samples, 50 *µ*L or homogenized body samples or 30 *µ*L saliva samples were tested by plaque assay, as previously described (121). The infection rate was defined as the ratio of plaque-positive body samples to the total number of mosquitoes (121). The transmission rate was defined as the ratio of plaque-positive saliva samples to the total number of mosquitoes (121).

For the longevity studies, post-segregation, mosquitoes were maintained on either 10% sucrose or water ad libitum at 28°C, 12:12 (light:dark) photocycle, and 75% relative humidity. Mosquitoes were assessed each day for survival, as determined by the number of live mosquitoes in the cage, and fecundity, as determined by the presence or absence of eggs. We used the hazard ratio (HR), which represents the relative risk of death between two groups, to compare survival probabilities. An HR greater than 1 indicates a higher risk of death in the group of interest compared to the reference group, which in this case is the colony-derived females.

To assess the extrinsic incubation period (EIP), mosquitoes were maintained on 10% sucrose *at libitum* using either cotton balls (days 2, 4, and 6 post-blood feed) or filter paper (days 1, 3, 5, 7, 8, 9, and 10 post-blood feed). Collected filter papers were stored in 100 *µ*L of mosquito diluent to elute virus from the filter papers (122). The eluent was assessed for the presence of virus using the NEB Luna^®^ Universal One-Step RT-qPCR Kit (E3005S). The forward primer was 5’-TTGGTCATGATACTGCTGATTGC-3’ and the reverse primer 5’-CCTTCCACAAAGTCCCTATTGC -3’ (123).

### H. Mathematical models of ZIKV transmission

Details of mathematical models including equations, parameter values, and model assumptions can be found in SI-2.

#### H.1. Larval Mass Model (LMM)

We develop a detailed model of mosquito population dynamics including egg, larval, pupal and adult stages that are separated by three distinct mass groups (small, medium and large). The overall structure of the mosquito population dynamics of the model is shown in Fig. 6A. The mosquito population model is incorporated into a transmission model including susceptible, exposed, infectious, and recovered humans (Fig. 6C). We split the human population into two groups: one that receives 90% of the mosquito bites and the other that receives 10% of the mosquito bites, as it is known that exposure to mosquitoes is heterogeneous (124). We assume that biting, and thus the potential transfer of the pathogen, occurs during the host-seeking stage of the gonotrophic cycle. As we do not include demographics or waning immunity, the human model of infection can only produce a single epidemic wave. The overall structure of the transmission model can be seen in Fig. 6C. The LMM is parameterized with experimental data as described in Section SI2. For parameters related to Zika infection in humans, we use literature derived values as described in SI-2.5.

#### H.2. Ross-Macdonald Model (RMM)

We use a modified Ross-Macdonald model (88, 125) that incorporates susceptible, exposed, infectious, and recovered humans and susceptible, exposed and infectious mosquitoes. For comparison to the LMM, we split each compartment of the human population into two groups based on heterogeneous biting. Parameterization for the RMM are taken to be equivalent to average parameterization from the LMM as described in Section SI-3.7.

## Supplementary Information

### SI-1. Transcriptomic Analysis

To investigate body size-dependent gene expression in *Ae. aegypti*, we conducted high-throughput RNA sequencing of whole-head tissues from large and small female mosquitoes. In mated females, sequencing yielded 14,909 transcripts out of the 19,585 annotated genes in the *Ae. aegypti* genome, among which 9674 transcripts met the threshold for reliable expression (FPKM > 1). Of these, 205 and 249 transcripts were uniquely expressed in large and small females, respectively, while 9,220 transcripts were co-expressed between the two groups (Fig. 4A2). Differential expression analysis identified 278 transcripts that significantly varied between size classes, including 126 enriched in large females and 152 enriched in small females. Similar patterns were observed in virgin females, where 9,903 transcripts were reliably detected. Of these, 310 and 225 transcripts were uniquely expressed in large and small females, respectively, while 9,368 were co-expressed across size groups. These findings underscore that while a large portion of the mosquito head transcriptome is shared across size classes, distinct subsets exhibit robust size-dependent expression.

Sequencing quality was consistently high across all libraries. For mated females, an average of 92.66% of reads from large females (range: 92.36–93.09%) and 92.68% from small females (range: 92.02–92.93%) aligned to the *Ae. aegypti* reference genome. Multi-mapping rates averaged 15.47% (range: 14.10–16.31%) in large females and 13.80% (range: 13.43–14.70%) in small females. For virgin females, sequencing quality was comparably high. An average of 92.90% of reads from large females (range: 92.44–93.32%) and 92.73% from small females (range: 92.05–93.15%) successfully aligned to the *Ae. aegypti* reference genome. Multi-mapping rates averaged 13.04% in large females (range: 12.61–13.68%) and 13.95% in small females (range: 11.96–17.74%). These values surpass standard RNA-seq quality thresholds, indicating excellent sequencing performance. To evaluate transcriptome coverage, we benchmarked our expression data from mated females against established standards, given that sequencing depth, library preparation, and mapping protocols were identical across all samples. Notably, 94 of the 100 least abundant genes identified by Hixson et al. (35) were reliably detected in our libraries, ruling out limitations due to shallow sequencing depth.

Moreover, 429 out of 451 annotated Core Eukaryotic Genes (CEGs)–a universally conserved set of housekeeping genes– were expressed above 1 FPKM in both large and small females, further validating the completeness and representativeness of transcriptome coverage across our dataset. Principal component analysis (PCA) of gene expression across biological replicates (n = 4 per group) revealed tight clustering within size classes, demonstrating strong intra-group consistency and validating the reproducibility of our sampling and library preparation pipeline. Although biological variation was modestly higher among large females, pairwise correlations between replicates consistently exceeded Pearson’s R^2^ > 0.942 across all comparisons (Fig. S4A,C), underscoring the robustness of transcript quantification. PCA also revealed clear separation between small and large females along the first two principal components, which accounted for 35.73% and 19.38% of total transcriptomic variance in mated females, and 31.12% and 18.43% in virgin females (Fig. S4B,D). These results establish adult body size as a dominant axis of transcriptional variation in female mosquitoes and provide a solid foundation for downstream analyses linking molecular signatures to size-dependent phenotypes.

### SI-2. Larval Mass Model (LMM)

Our larval mass model (LMM) is a detailed model of mosquito population dynamics including egg, larval, pupal and adult stages and is an extension of previous work (126). We track three distinct mass groups (small, medium and large). As larvae, individuals grow (up to a max ‘large’ size) and thus move up to higher mass groups. Once an individual becomes a pupa, growth is no longer considered, although individuals are still categorized by mass. Once a female mosquito emerges as an adult, she is considered a young adult, denoted by *Y*, as she does not immediately enter a gonotrophic cycle to find a host. Once the female enters the gonotrophic cycle there are three stages denoted by *H, R*, and *L* for host-seeking, resting, and egg laying, respectively. This process is repeated for eight gonotrophic cycles, after which we assume all remaining mosquitoes die. During all adult stages, there is natural mosquito mortality as described in Section SI-3.1. Males are only explicitly modeled in aquatic stages. For equations of the LMM, see Section SI-2.1 with details on the parameterization in Section SI-3. A summary of variables and parameters for the LMM are given in Tables S1 and S2, respectively.

#### SI-2.1. Mosquito Population Dynamic Equations

In previous work (126), we began at the first larval (*L*1) stage, but now we include eggs. No eggs are discarded as we fit all birth (*i*.*e*., egg hatching) functions *β*(*s*), *β*(*m*), and *β*(*𝓁*) to be the total progeny. As eggs may not immediately hatch (127), in our model we force them to take a minimum of two days to hatch into *L*1. The proportion of eggs that hatch each day is fit from data as described in Section SI-3.2. While all aquatic equations are described in detail in previous work (126), we have included the equations below for completeness. As we are unable to sex the individuals until emergence as adults, we assume that, in each replicate, 50% of the initial larvae are male, and 50% are female. Individuals develop through successive larval stages, where the proportion of juvenile mosquitoes that develops each day is governed by the function *f* (*N*). At each transition, a proportion of individuals die *µ*(*N*) as a function of the total larval population. For our discrete time model, we assume that at the end of each period that mosquitoes transition first and then introduce mortality.

During the larval stages, mosquitoes may remain in their current mass group or transition to a higher mass group with the assumption that mosquitoes changing mass groups can only transition to the next highest mass group in a single time step. For example, during the transition from the first larval stage (*L*1) to the second larval (*L*2) stage, individuals can either stay in the small mass group or move to the medium mass group. Similarly, each individual in *L*2 that transitions to the third larval stage (*L*3) can stay in the same mass group or move to a higher mass group.

Note that for the larval and pupal stages, we track males and females separately to account for differences in development and survival that are sex-dependent, but we only present larval and pupal equations for a single sex here as the equations are structurally identical. While the equation structure is identical, the following functions differ depending on sex: death proportion, *µ*(*N*); the proportion which develop, *f* (*N*); and the proportion which grow, *G*_1_(*N*), *G*_2_(*N*), where *N* is total larvae (males + females). Details of these functions is found in (126).

Equations for the egg, larval and pupal compartments are given by

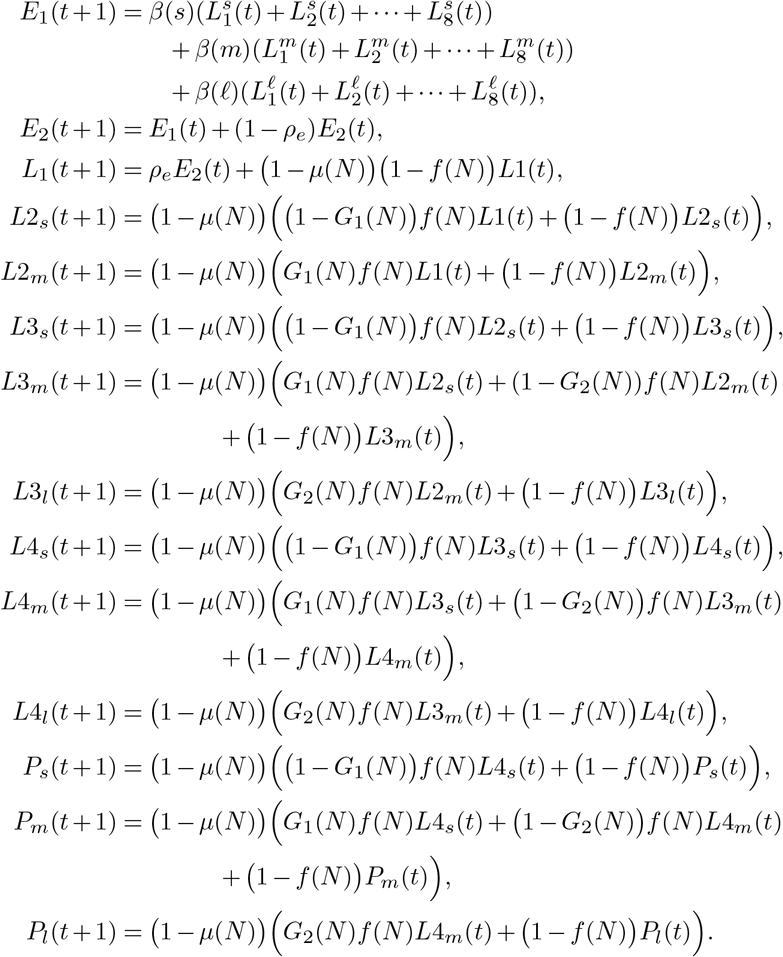

In our model, we denote the second compartment in each of *L*4 and pupal stages with an asterisk. For example, *P*_*l*_ is the first compartment for pupae in the large mass group and 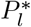 is the second compartment. No growth is assumed during these transitions so they take the form

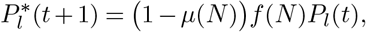

which is from (126).

Adult females are denoted by stage, so that *Y, H, R*, and *L* represents pre-gonotrophic stages, host-seeking, resting, and egg laying, respectively. For simplicity, we exclude notation for each mass group, but all equations are repeated for all three mass groups (small, medium, large). The only difference between the equations by mass are the associated parameters.

There is a delay before females host-seek ranging from 5 to 13 days, so pre-gonotrophic cycle boxes are split into several substages and denoted ordinally by subscript. For example, *Y*_2_ represents pre-gonotrophic individuals on the second day after emergence. Starting on day five, some individuals move to host-seeking for the first gonotrophic cycle (*H*_1_) and the rest move to *Y*_6_, with more moving to *H*_1_ each day. By the 13th day all mosquitoes move to *H*_1_. This is fitted from the data based on small or not small mosquitoes as described in Section SI-3.4.

After adult females reach host-seeking compartments (*H*_1_, *H*_2_, *…, H*_8_), a proportion (*h*_*day*_(mass group)) move to the rest compartment. All individuals stay exactly one day in the respective rest compartment (*R*_*ja*_, *R*_*jb*_, *…, R*_*jd*_) for each gonotrophic cycle *j*. However, the number of rest days differ based on mass, and the fitting is described in Section SI-3.3. Adult females move to egg laying compartment after three to five days of rest. For each cycle, there is a single egg laying compartment, in which females lay eggs and spend exactly one day before moving on.

Equations for the young adult compartments, where *i* refers to mass group, are given by

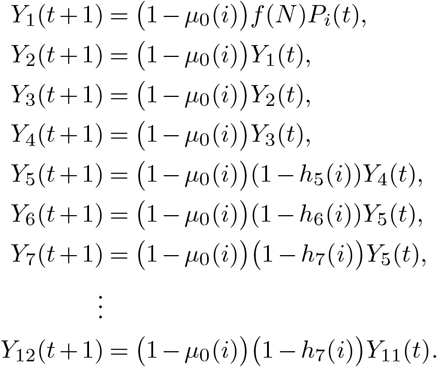

Equations for the gonotrophic cycle adult compartments are given by:

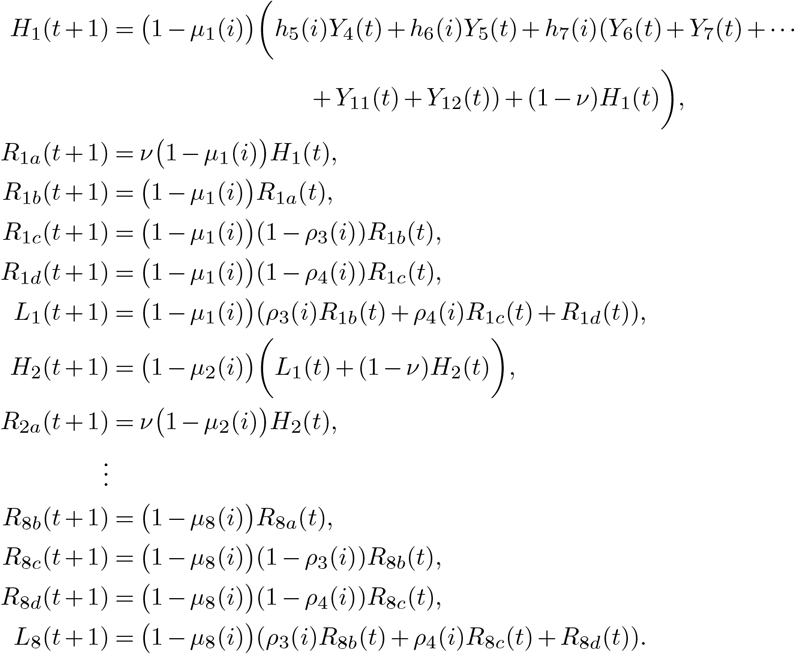

As mentioned above, these equations are repeated for each mass group (small, medium, and large), and only the parameterization differs.

#### SI-2.2. Zika transmission

We extend the model of mosquito population dynamics to include transmission of the mosquito-borne pathogen Zika. To do so, we include equations for exposed and infectious mosquitoes as well as for susceptible, exposed, infectious and recovered humans.

The act of blood feeding can transfer pathogen from an infectious mosquito to a human or from an infectious human to a mosquito. Thus, we assume that a susceptible mosquito can become infected or an infectious mosquito can infect when the mosquito moves from host-seeking to resting as they have ingested a bloodmeal. When a susceptible mosquito bites an infectious human, the mosquito is exposed to Zika with probability *α*_*v*_. If the mosquito becomes infected, it moves into the exposed class for one full gonotrophic cycle. On the second gonotrophic cycle after acquiring the pathogen, the mosquito is infectious to humans. Due to the discrete nature of the model, this forces a minimum incubation time for Zika in mosquitoes of six days. To account for the incubation period, we separate mosquitoes within the exposed class who are infected at the first bite from those infected later.

If an infected mosquito bites a susceptible human, the probability of infection is *α*_*h*_. Once a human receives an infectious bite from a mosquito, the individual moves to the exposed class. A daily proportion of infected humans, given by *γ*_*E*_, move to the infectious class. Following infection, a daily proportion of humans, given by *γ*_*I*_, recover after which they remain immune. As we do not include demographics or waning immunity, the human model of infection can only produce a single epidemic wave.

#### SI-2.3. Force of Infection

In order to determine the force of infection for mosquitoes (i.e., human to mosquito transmission) given by *λ*_*v*_, we consider the proportion of the human population that is infectious. As it is known that exposure to mosquitoes is heterogeneous (124), we split the human population into two groups: one that receives 90% (*N*_2_) of the mosquito bites and the other that receives 10% (*N*_1_) of the mosquito bites. We then multiple this fraction by the proportion of mosquitoes that successfully bite a human each day, *ν* and the probability that an infectious bite leads to infection, *α*_*v*_. Thus, the force of infectious for mosquitoes, which is also the proportion of mosquitoes that receives an infectious bite, is

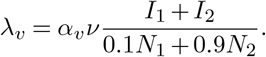

For the force of infection for humans (i.e., mosquito to human transmission) given by *λ*_*h*_, we must include all infectious mosquitoes that successfully bite a human each day, *ν*. We sum all bites, scaled by probability that a bite from a particularly sized mosquito will cause human infection, *α*_*hk*_ with *k* ∈{*s, m, l*}, and divide by the human population weighted by propensity for being bitten. Thus, the force of infection for humans is

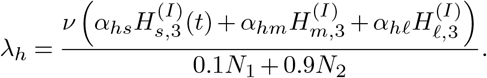

If the human population is smaller than the number of bites on a given day, both force of infection equations have the potential to be greater than one, which does not make sense mathematically. To ensure that this does not occur, we restrict both *λ*_*v*_, *λ*_*h*_ ≤ 1.

#### SI-2.4. Equations for Zika Transmission

As with the LMM without infection, the mosquito equations are repeated for each mass group: small, medium, and large. The equations for the susceptible mosquitoes are given by

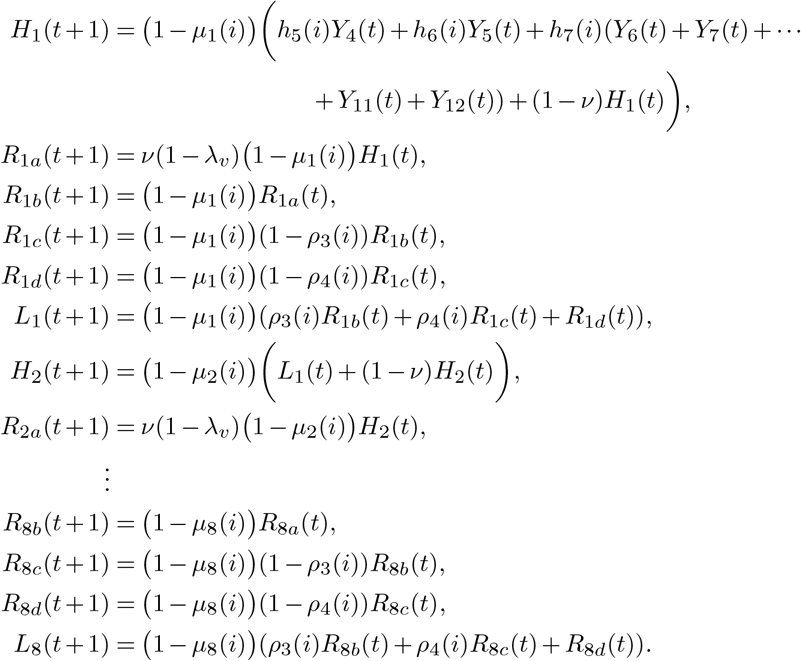

These are very similar to the original model except at the host-seeking to resting transition, individuals can become infected and move to the exposed system of equations.

The equations for exposed mosquitoes are given by:

Exposed after first cycle

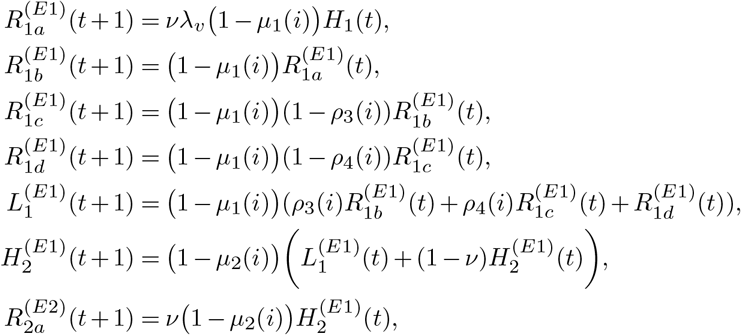

Exposed after second cycle

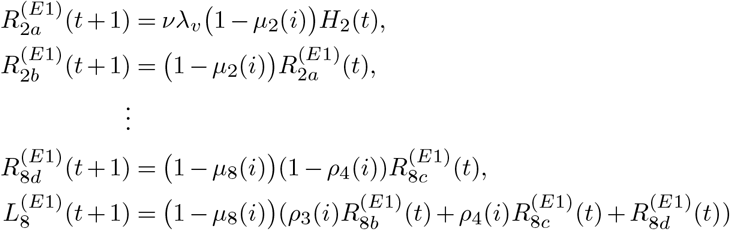

The equations for infectious mosquitoes are given by

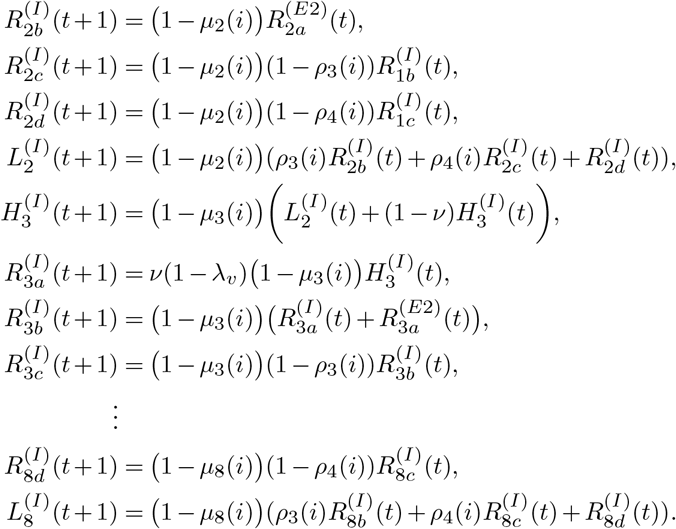

The equations for human compartments are given by

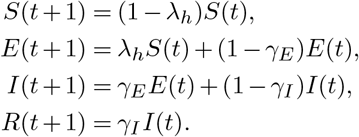

As discussed in the previous section, the force of infection for mosquitoes and humans is given by *λ*_*v*_ and *λ*_*h*_, respectively.

### SI-3. LMM Parameterization

#### SI-3.1. Death proportion by mass group and stage

To fit separate death proportions for each mass group, we first determine that each mass group survival is statistically significant (Fig. S6A, log-rank test p = 0.021). We use data from two different experiments: in the first experiment, mosquitoes were only given water, while in the second, the mosquitoes were given sugar as well as water.

In order to fit the data well, we specify eight specific gonotrophic cycles. For each mosquito mass group, we estimate three parameters (i.e., three survival proportions), which cover the periods of the pre-gonotrophic cycle stages as well as all stages in the eight gonotrophic cycles. This choice to fit three parameters is motivated by the survival probability curves in Fig. S6A, in which there are three distinct changes in survival. For all mass groups, there is very little initial decrease in survival followed by a sharp decline and, finally, a very shallow decline such that older mosquitoes which persist tend to have extended survival. We fit one survival proportion for each of these three distinct phases of the survival curves, whose timing differs by mass. For small mosquitoes the mosquito survival drops early, so we fit an initial survival proportion for the pre-gonotrophic cycle stages, a second proportion for the first four gonotrophic cycles, and a third proportion for the last four gonotrophic cycles. For medium and large mosquitoes, the pre-gonotrophic cycle stages and the first gonotrophic cycles show the same survival trend so we fit these with the first survival proportion. We fit a second proportion for the second, third and fourth gonotrophic cycles and a third proportion for the last four gonotrophic cycles.

#### SI-3.2. Hatch distribution

For all egg batches, eggs hatch at varied times, but there is no difference in the timing of hatching for each mass group (Fig. S6C). To determine the percentage of eggs that hatch each day, we fit the data of eggs that hatch per day for each female to a geometric distribution. Instead of considering viability and hatch time separately, we only fit the daily hatch probability from eggs that hatch, and assume the number of eggs in the batch that die or are not viable, die immediately. There is a significant difference between the number of eggs laid by mass group (Kruskal-Wallis Rank Test *p* = 0.004, Welch’s ANOVA *p* = 2.5 *·* 10^*−*5^).

#### SI-3.3. Gonotrophic cycle length

For each gonotrophic cycle, the proportion of mosquitoes that laid eggs on a particular day is used directly in the model, measured in days post blood feed and categorized by size (Fig. S6B). While no mosquitoes laid their first eggs later than six days after blood feeding, due to the discrete nature of the model, we cut attempts at egg laying off at five days. Therefore, data on medium mosquitoes that laid on day six are included in the proportion that lay on day five. Fisher’s exact test (*p* = 6.9 *·* 10^*−*6^) indicates that the daily laying proportions differ significantly across size classes.

#### SI-3.4. First gonotrophic cycle length

The first gonotrophic cycle takes longer since mosquitoes must reach maturity before they start blood feeding. To account for this, the measured time since emergence to first blood feeding was used for the length of the first gonotrophic cycle. When grouping by size, the difference in the length of the first gonotrophic cycle was only significant (based on a log-rank test p = 0.04) when large and medium mosquitoes were combined (Fig. S6D).

The measured timing on which mosquitoes take their first blood meal following emergence ranged from day five to day twelve. For days five and six, we use the proportion of mosquitoes that blood feed directly from the experimental data, but for days seven through twelve we fit a single proportion per day (using a least squares error function) to the actual proportion which moves to host-seeking in the experimental data (Fig. S6E). In the model, all mosquitoes who have not blood fed or died after twelve days move to blood feeding on day 13.

#### SI-3.5. Transmission model parameterization

The probability of mosquito infection (*α*_*v*_) is determined from the proportion of mosquitoes where Zika was found in the body following infection (Fig. 55G). The probability of infection of humans by mosquito size (*α*_*hj*_ with *j* ∈ {*s, m, l*}) is determined from the proportion of mosquitoes where Zika was found in the saliva following infection (Fig. 5H). For parameterization related to Zika infection in humans, we use literature derived values. We assume the average latent period is four days (128–130) and the average infectious period is four days (128, 129, 131). Thus, *γ*_*E*_ = *γ*_*I*_ = 1 *−* exp(−4).

To determine scaling between mosquito and human populations, we fit the concentration of humans so the total cumulative infections (as a proportion of the total population) in a modeled outbreak matches the total outbreak proportion in the Yap Island outbreak with all other parameters fixed. In the Yap Island data 73% of the human population was estimated as infected during the epidemic (132), so our standard outbreak is 73% of the population, which we use as the reference value for all interventions. This yields a human concentration of 2.36 in standard LMM conditions.

### Ross-Macdonald Model (RMM)

For comparison to the LMM, we construct a simple Ross-Macdonald based model (RMM).

#### SI-3.6. RMM Equations

The RMM has susceptible, exposed and infectious mosquitoes and susceptible, exposed, infectious and recovered humans. For consistency, we split the human compartments into two for each stage to account for heterogeneous biting (124) with weight *ω*_*i*_ for type *i*. The force of infection includes biting probability (*a*) and probability of infections *α*_*v*_ and *α*_*h*_ for mosquitoes and humans, respectively. Transitions from exposed to infectious occur with probability *σ*_*k*_ for *k* ∈ {*h, v*} and recover of humans with probability *γ*_*hI*_. Mosquitoes are recruited with probability Λ_*v*_ and die with probability *µ*_*h*_. As with the LMM model, we assume no human demographics or waning immunity. All variables for the RMM are summarized in Table S3.

The mosquito equations are given by

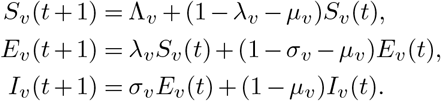

The human equations for type *j* for *j ∈* {1, 2} are given by

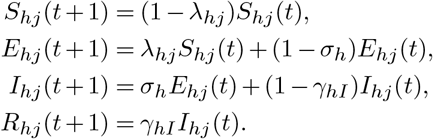

In the above equations, the forces of infection are given by

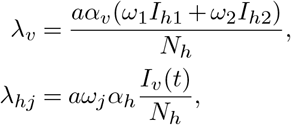

where the total human population is given by

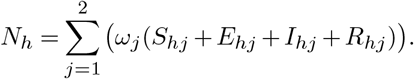

#### SI-3.7. RMM Parameterization

Parameters for the RMM are chosen to be as consistent as possible with those in the LMM and are summarized in Table S4. As with the LMM, multiple parameters were taken from the data in the present study. The probability of mosquito infection (*α*_*v*_) is determined from the proportion of mosquitoes where Zika was found in the body following infection averaged across mass groups (Fig. 5G). The probability of infection of humans by mosquito (*α*_*h*_) is determined from the proportion of mosquitoes where Zika was found in the saliva following infection averaged across mass groups (Fig. 5H). The average lifespan of mosquitoes in the experimental data was 20.5 days so we assume that Λ_*v*_ = *µ*_*v*_ = 1 − exp(−1*/*20.5). From the data, there were approximately 0.15 bites per day such that *a* = 1 −exp(−0.15). For parameterization related to Zika infection in humans, we use literature derived values as for the LMM. We assume that *σ*_*h*_ = *γ*_*E*_ = 1 −exp(−1*/*4) and *γ*_*h*_ = *γ*_*I*_ = 1 −exp(−1*/*4) (128–131). The latent period of mosquitoes is chosen to be 10 days such that *σ*_*v*_ = 1 −exp(−1*/*10) (129, 133, 134). The human to mosquito ratio was chosen to be between 2.36 and 33 for consistency with the LMM.

## Supplementary Figures

**Fig. S1.**
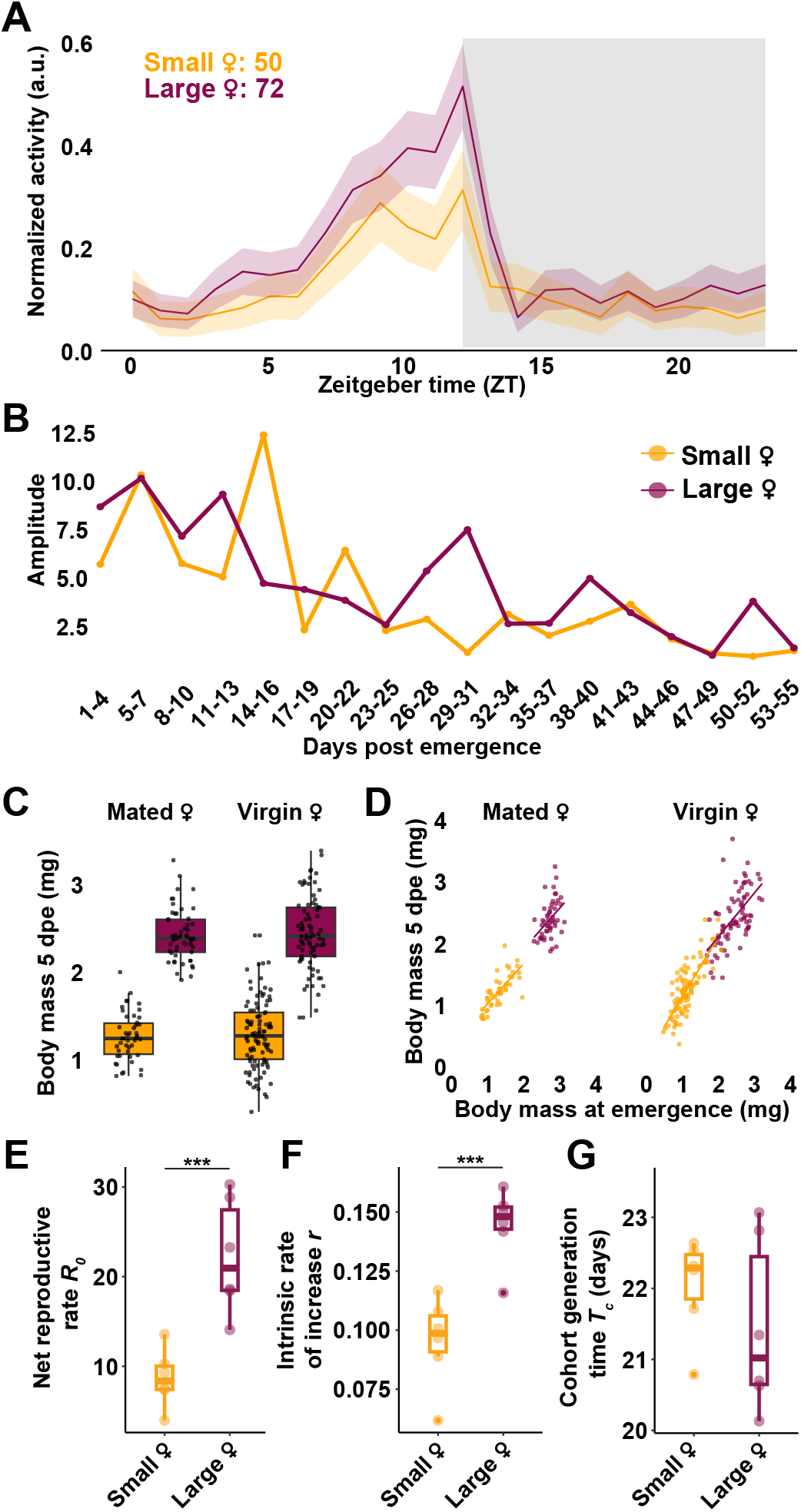
Locomotor activity rhythms and body mass dynamics in small and large female mosquitoes. **(A)** Normalized 24-hour locomotor activity profiles of small (orange, n = 50) and large (maroon, n = 72) mated females recorded under a 12:12 h light:dark cycle. Shading in grey denotes the dark phase. **(B)** Locomotor rhythm amplitude plotted across consecutive 3-day intervals through mosquitoes’ lifespan post-emergence. **(C)** Body mass at 5 days post-emergence (dpe) in mated and virgin females reared under high- and low-density larval conditions. **(D)** Body mass at 5 dpe plotted as a function of body mass at emergence for mated and virgin females. Each point represents an individual female. **(E–G)** Life table parameters of small and large female mosquitoes: **(E)** Net reproductive rate (*R*_0_), **(F)** Intrinsic rate of increase (*r*), and **(G)** Cohort generation time (*T*_*C*_) calculated for small and large females using age-specific survival and fecundity data. Each point represents an independent replicate cohort. Statistical comparisons were made using ANOVA.

**Fig. S2.**
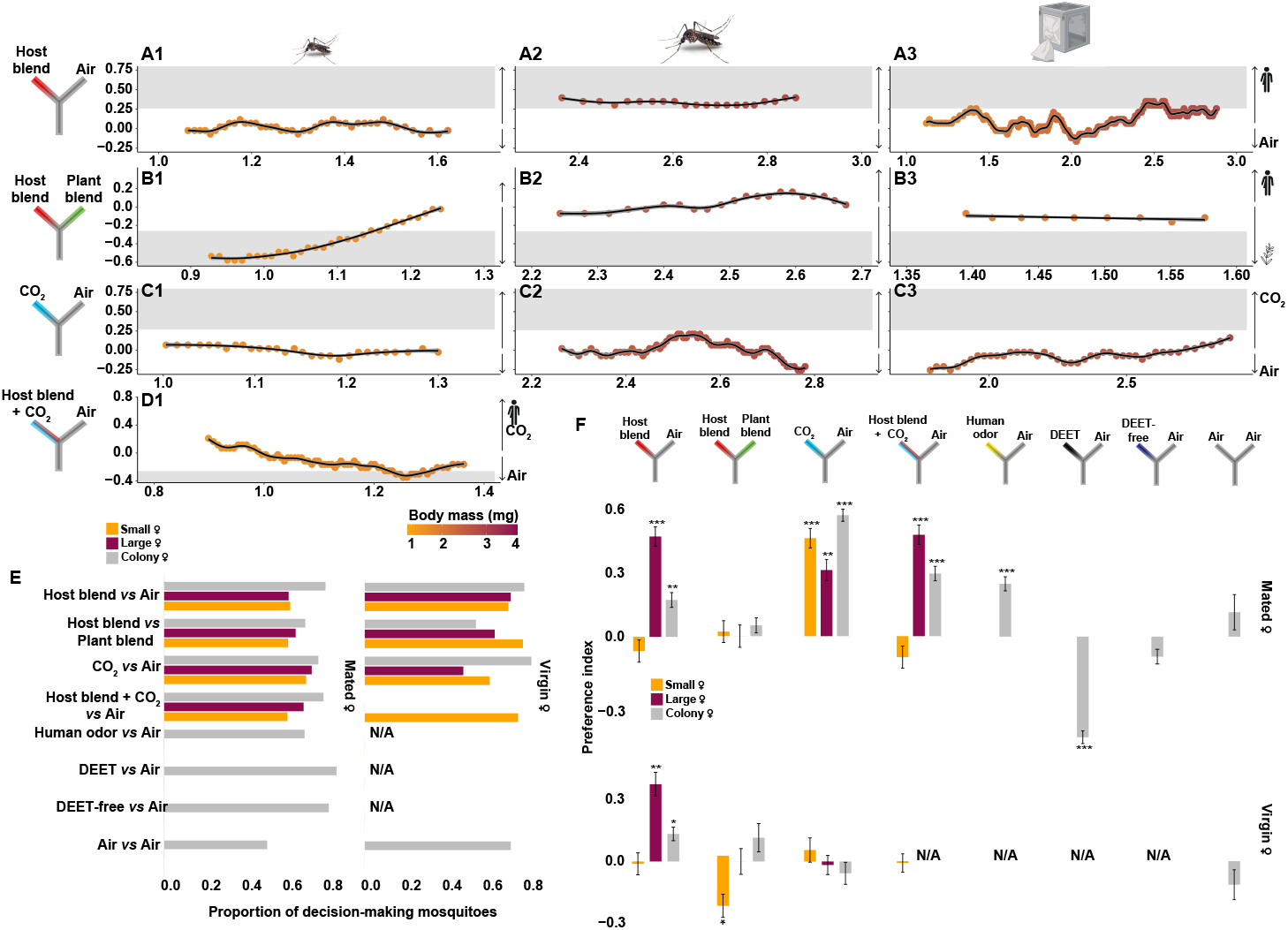
Mating status and body size shape mosquito olfactory preferences for host odors, plant odors, and CO_2_. Olfactory preferences were quantified using Y-maze olfactometer assays across a range of odor stimuli and mosquito body sizes. **(A1–D1)** Small virgin females reared at high larval density, **(A2–C2)** large virgin females reared at low larval density, and **(A3–C3)** colony-derived, virgin females representing the full range of adult body sizes were tested. Odor contrasts included: **(A1–A3)** host blend vs. clean air, **(B1–B3)** host blend vs. plant blend, **(C1–C3)** CO_2_ vs. clean air, and **(D1)** host blend + CO_2_ vs. clean air. Preference index is plotted as a function of individual body mass (mg). **(E)** Proportion of decision-making mosquitoes in each assay, shown separately for small, large, and colony-derived females across odor contrasts and reproductive states - mated (left panel) and virgin (right panel). **(F)** Mean preference index (PI) values for small (orange), large (maroon), and colony (gray) females, mated (top panel) and virgin (bottom panel), across odor contrasts, with responses to host blend, plant blend, CO_2_, host blend + CO_2_, human odor, DEET, and DEET-free plant-derived repellent, each tested against clean air. The preference index (PI) represents the difference in proportions choosing the odor stimulus versus the control, ranging from –1 (complete avoidance) to +1 (complete attraction). Data points overlapping with the shaded region in **(A–D)** indicate statistically significant preference indices. Statistical significance was assessed via one-tailed Binomial Exact tests; trend lines were modeled using a Generalized Additive Model. Statistical significance is indicated by asterisks (** *p* < 0.01; *** *p* < 0.001). Error bars denote mean ± s.e.m. Exact sample sizes and summary statistics are provided as source data.

**Fig. S3.**
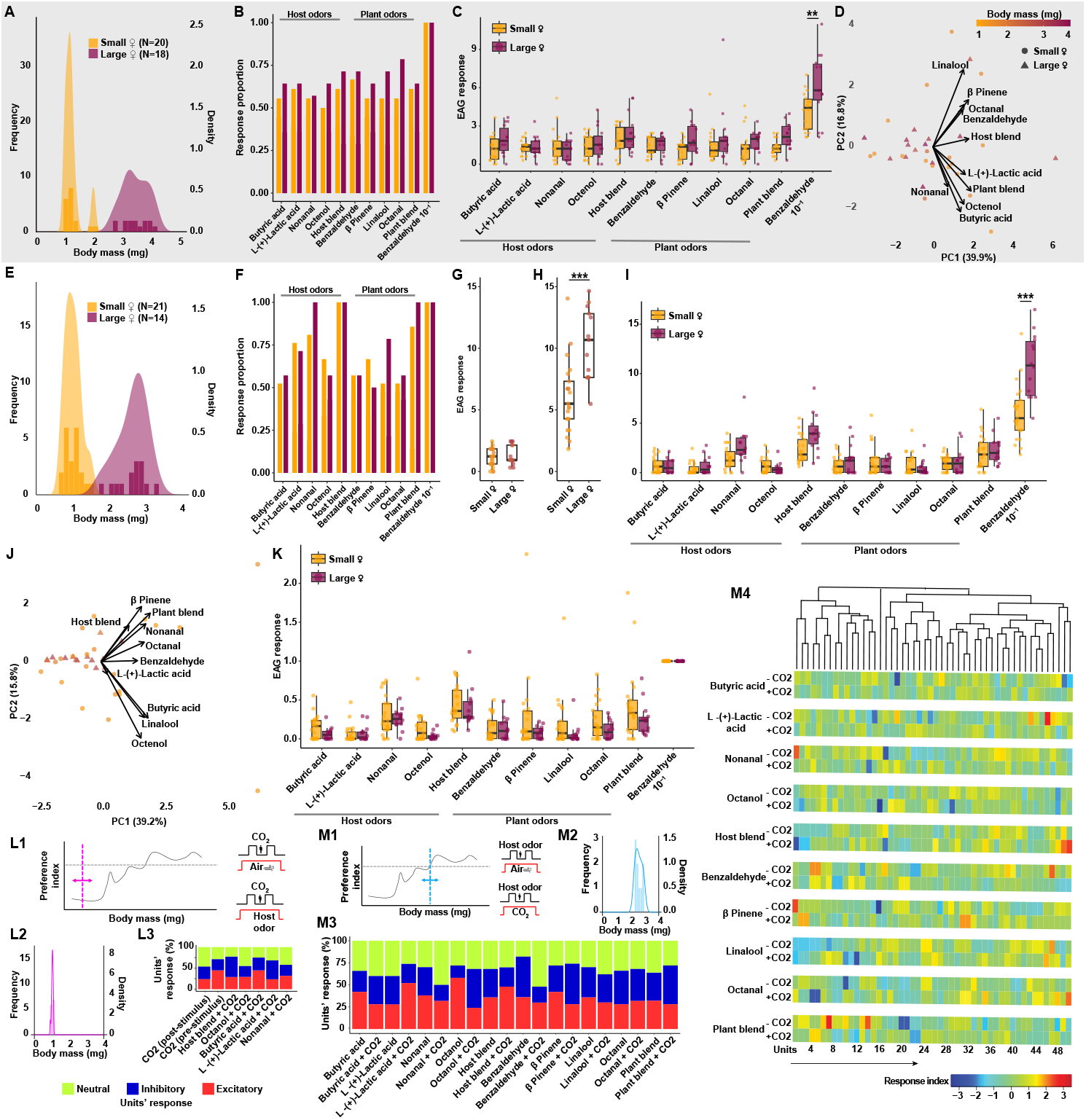
Peripheral olfactory sensitivity is body size-independent across mating states, but CO_2_ modulates central odor encoding in the mosquito brain. **(A–D)** Electroantennogram (EAG) responses of small and large mated females (shaded in grey) to host and plant odorants. **(A)** Body mass distributions of small (orange) and large (maroon) mated females subjected to the analysis. **(B)** Proportion of individuals showing detectable responses to host and plant odorants. **(C)** EAG amplitudes evoked by individual odorants in the host and plant blends. Each data point represents an individual mosquito’s response, normalized to mineral oil. **(D)** Principal component analysis (PCA) of odor-evoked EAG responses with body mass variations represented by color. **(E–K)** EAG responses of small and large virgin females. **(E)** Body mass distributions of small (orange) and large (maroon) virgin females subjected to the analysis. **(F)** Proportion of individuals showing detectable responses to host and plant odorants. **(G)** Response to mineral oil (negative control). (H) Response to benzaldehyde (positive control, 10^*−*1^ concentration) normalized to the negative control. **(I)** EAG amplitudes evoked by individual odorants in the host and plant blends. Each data point represents an individual mosquito’s response, normalized to mineral oil. **(J)** Principal component analysis (PCA) of odor-evoked EAG responses with body mass variations represented by color. **(K)** Responses to the host odor blend and its constituents (butyric acid, L-(+)-lactic acid, nonanal, and octenol), the plant odor blend and its constituents (benzaldehyde, *β*-pinene, linalool, and octanal), and the positive control (10^*−*1^ concentration). Each data point represents an individual mosquito’s response normalized to mineral oil and visualized relative to the response to positive control. Statistical significance (in panels C,G-I, and K) was determined using a two-factor ANOVA with Tukey post-hoc test and Bonferroni correction (*p* < 0.05). **(L1–L3)** Neural activity characterization for antennal lobe recordings. **(L1)** Schematic illustrating the selection of mated females for extracellular recordings. Left: body mass distribution with the selected range indicated by a vertical line and arrow. Right: schematic of stimulus delivery in the assay depicting CO_2_ pulses delivered against a background of either host odor blend or clean air. **(L2)** Body mass distribution of all mated females subjected to antennal lobe recordings under the conditions shown in L1. **(L3)** Response classification of neural units based on extracellular recordings, conducted under the conditions described in L1, showing proportions of excitatory, inhibitory, and neutral responses across odorants and conditions. **(M1–M4)** Neural responses to host and plant odorants in the presence or absence of CO_2_. **(M1)** Schematic illustrating the selection of mated females for extracellular recordings. Left: body mass distribution with the selected range indicated by a vertical line and arrow. Right: stimulus delivery setup depicting odor pulses (host or plant blend and constituents) presented on backgrounds of either CO_2_ or clean air. **(M2)** Body mass distribution of all mated females subjected to subjected to antennal lobe recordings under the conditions shown in M1. **(M3)** Response classification of neural units based on extracellular recordings, conducted under the conditions described in M1, showing proportions of excitatory, inhibitory, and neutral responses across odor stimuli and CO_2_ backgrounds. **(M4)** Neural ensemble response matrix depicting the activity of 50 neural units across 9 preparations in response to CO_2_ alone, CO_2_ + host blend, and CO_2_ with individual host blend constituents.

**Fig. S4.**
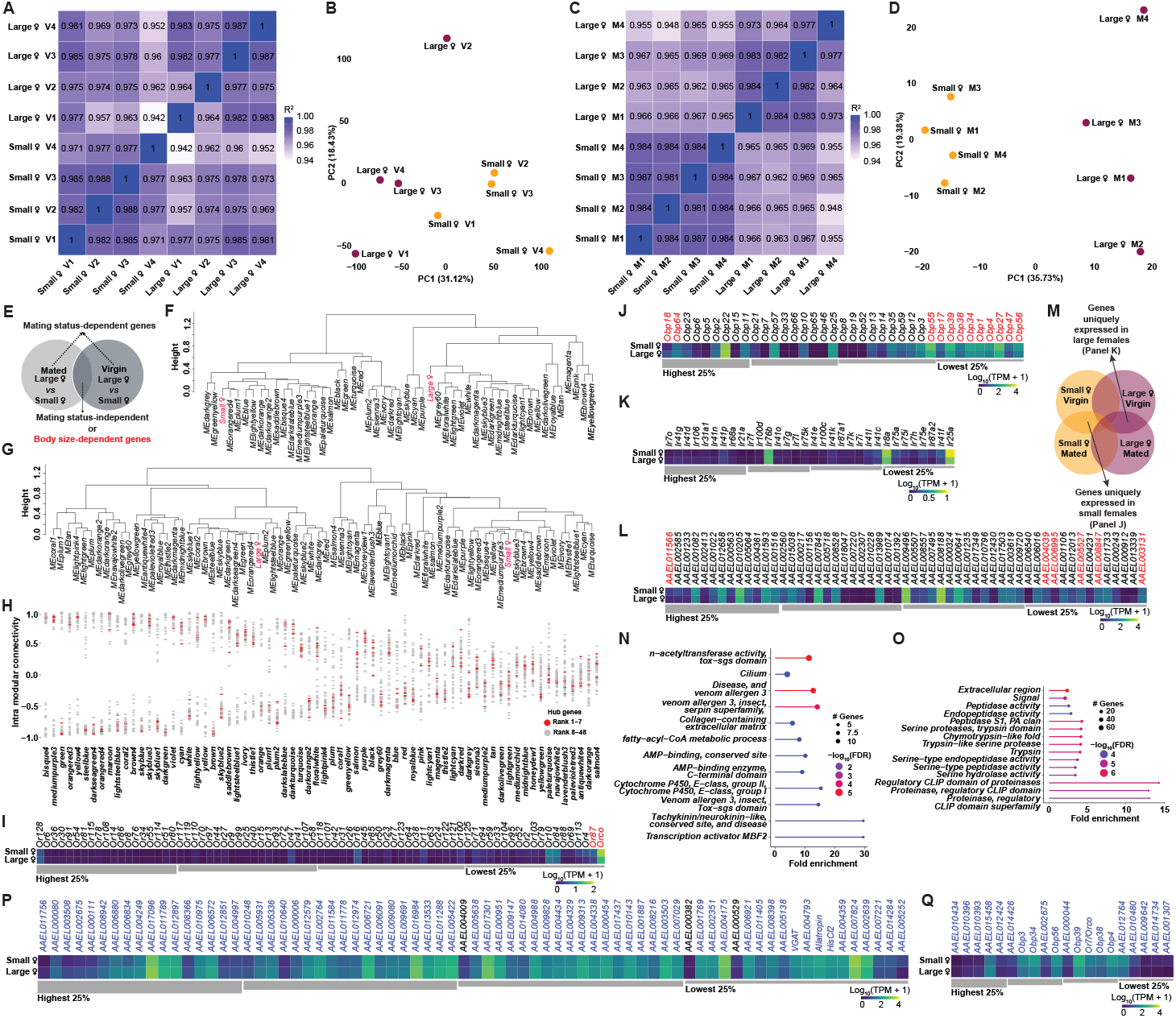
Integrated transcriptomic analyses reveal body size- and mating status-specific gene networks in female mosquitoes. **(A–D)** Quality assessment and global transcriptomic clustering of RNA-seq libraries from small and large females across mating states. **(A, C)** Correlation heatmaps for virgin **(A)** and mated **(C)** females show high within-group consistency (R^2^ > 0.96) and lower between-group correlations, indicating size-dependent expression patterns. **(B, D)** PCA of reliably expressed genes (FPKM > 1) reveals tight clustering of biological replicates by body size and clear separation between small and large females, both in virgin **(B)** and mated **(D)** groups, confirming robust size-associated transcriptomic divergence. **(E)** Schematic illustrating the experimental design used to disentangle the effects of body size and mating status on transcriptomic variation in whole-head tissues of female mosquitoes. **(F–G)** Eigengene dendrograms from WGCNA showing clustering of gene modules based on expression similarity in virgin **(F)** and mated **(G)** females. Small and large females are indicated in red in each dendrogram. Modules clustering closer to either phenotype represent gene networks whose expression patterns are more strongly associated with that body size class. These dendrograms highlight body size-associated transcriptomic architecture across mating conditions. **(H)** Intramodular connectivity plot from WGCNA analysis of mated females, highlighting the network topology of the 48 candidate genes identified in Fig. 4G. Genes are ranked by their connectivity within their respective co-expression modules, with red dots denoting the seven hub genes and gray dots representing the remaining 41 candidates (ranks 8–48). Hub genes exhibit the highest intramodular connectivity, indicating their centrality in the transcriptional architecture and reinforcing their likely regulatory influence over body size-associated gene networks. **(I-L)** Differential transcript abundance is presented for key gene categories in virgin females: odorant receptors **(I)**, odorant-binding proteins **(J)**, ionotrophic receptors **(K)**, and immune function-related genes **(L)**. Gene names in red denote transcripts that are significantly differentially expressed between size classes (adjusted *p* 0.05). **(M)** Schematic illustrating the experimental design used to identify body size-dependent gene expression among uniquely expressed transcripts in whole-head transcriptomes of female mosquitoes. This analysis complements the approach in panel E by isolating transcripts found exclusively in either small or large females. **(N–O)** Gene Ontology (GO) enrichment analysis of transcripts uniquely expressed in small **(N)** and large **(O)** female mosquitoes. **(P-Q)** Differential transcript abundance of genes identified through Protein-Protein Interaction (PPI) network analysis using CytoHubba and CytoNCA algorithms for virgin **(P)** and mated **(Q)** females. Panels depict key regulatory genes subjected to analysis in Fig. 4F. Gene names in blue indicate consensus genes identified by both CytoHubba and CytoNCA, while those in black were identified by CytoNCA alone.

**Fig. S5.**
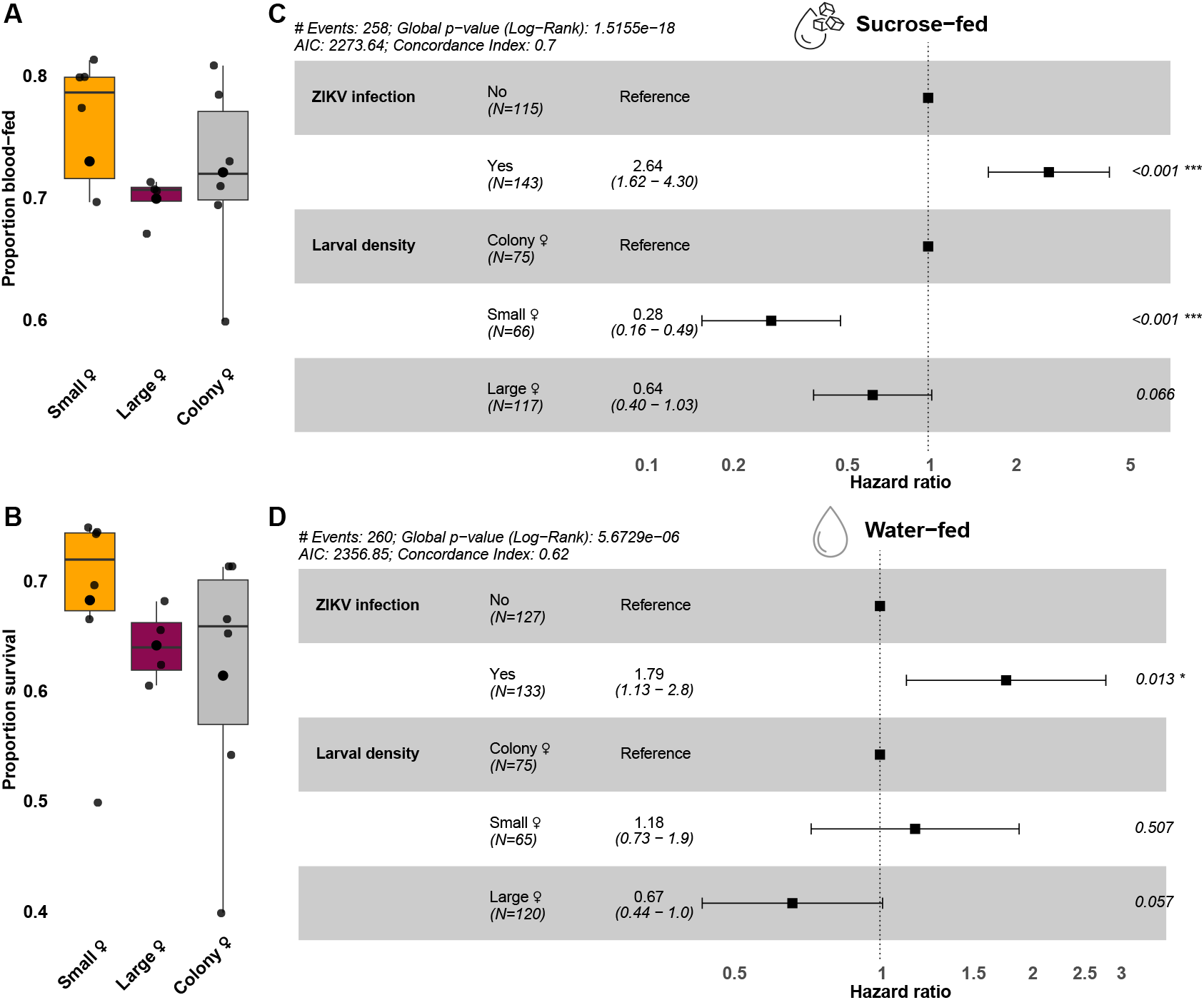
Influence of larval density, ZIKV infection, and post-blood meal diet on mosquito blood-feeding success and survival. **(A–B)** Proportion of females that **(A)** successfully ingested a ZIKV-infected blood meal and **(B)** survived over a 10-day period post blood feeding. These data correspond to the same cohorts of small, large, and colony-derived females whose infection and transmission potential are reported in Fig. 5G and 5H, respectively. Statistical significance between proportions was assessed using Generalized Linear Model (GLM) assuming quasibinomial errors. No significant differences were detected across treatment groups in **(A)** and **(B)** (*p* > 0.05). **(C–D)** Forest plots showing hazard ratios from Cox proportional hazards models assessing the effects of larval rearing conditions and ZIKV infection on adult mosquito survival when fed with 10% sucrose **(C)** or water **(D)** following the blood meal. Points represent estimated hazard ratios, with horizontal bars indicating 95% confidence intervals. Sample sizes **(N)** for each condition and 95% confidence intervals are shown alongside the corresponding hazard ratios. The dashed vertical line indicates the reference hazard ratio (HR = 1.0). Statistical significance is denoted by asterisks (* *p* < 0.05; *** *p* < 0.001). Model outputs and summary statistics are included on top of each figure.

**Fig. S6.**
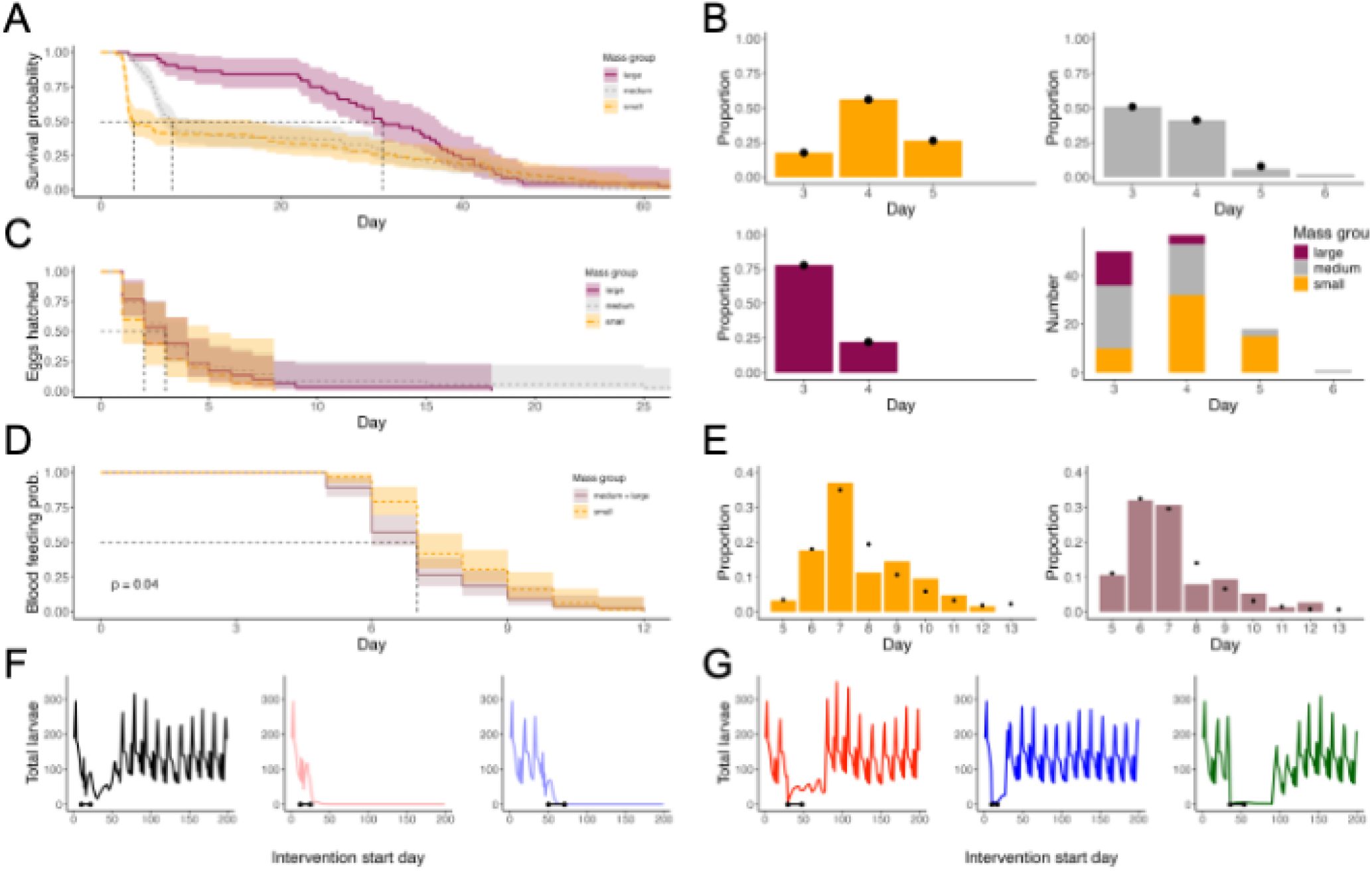
Model parameterization and additional results. **(A)** Kaplan Meier survival curves by mass group (shaded area indicates 95% confidence interval), where dashed black lines indicate the day at which 50% of mosquitoes remain. **(B)** Proportion of mosquitoes that lay eggs on a particular day post blood feeding by mass group: large mosquitoes in upper left; medium mosquitoes in upper right; small mosquitoes in lower left. Lower right shows the number of mosquitoes by mass group laying eggs by day. Colored bars are data and black dots are values used in the model. **(C)**Fraction of eggs hatched each day by mass group (shaded area indicates 95% confidence interval), where dashed black lines indicate the day at which 50% of eggs remain unhatched. 10^*−*1^ Fraction of mosquitoes who have taken their first blood feeding each day by mass group (shaded area indicates 95% confidence interval), where dashed black lines indicate the day at which 50% of mosquitoes have fed. Note that medium and large mosquitoes are combined. **(E)** Proportion of mosquitoes by mass group that take their first blood meal post emergence on each day (left: small mosquitoes; right: medium and large mosquitoes combined in one group). Colored bars are data and black dots are values used in the model which were obtained from model fitting. **(F)** Time series plots of total larvae showing different start time and durations for adult interventions (left to right: 90% starting on day 10 with a duration of 12 days; 95% starting on day 12 with a duration of 13 days; 99% starting on day 40 with a duration of 21 days). **(G)** Time series plots of total larvae showing different start time and durations for larval interventions (left to right: 90% starting on day 30 with a duration of 18 days; 95% starting on day 10 with a duration of 7 days; 99% starting on day 36 with a duration of 18 days). Time series plots in (F) correspond to results in Fig. 6E,G and those in (G) correspond to results in Fig. 6F,H.

## Supplementary tables

**Table S1.**
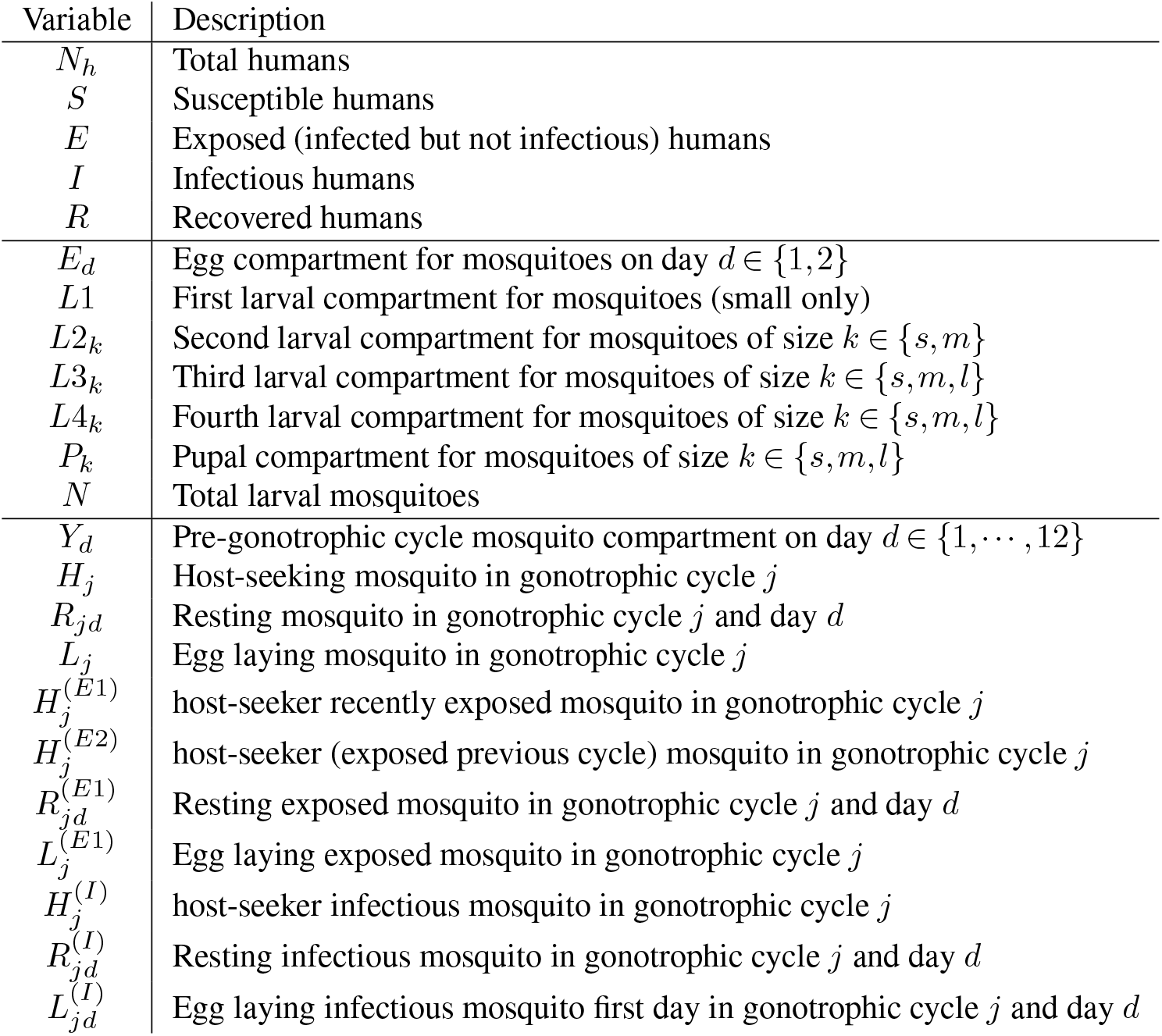
Variables in LMM.

**Table S2.**
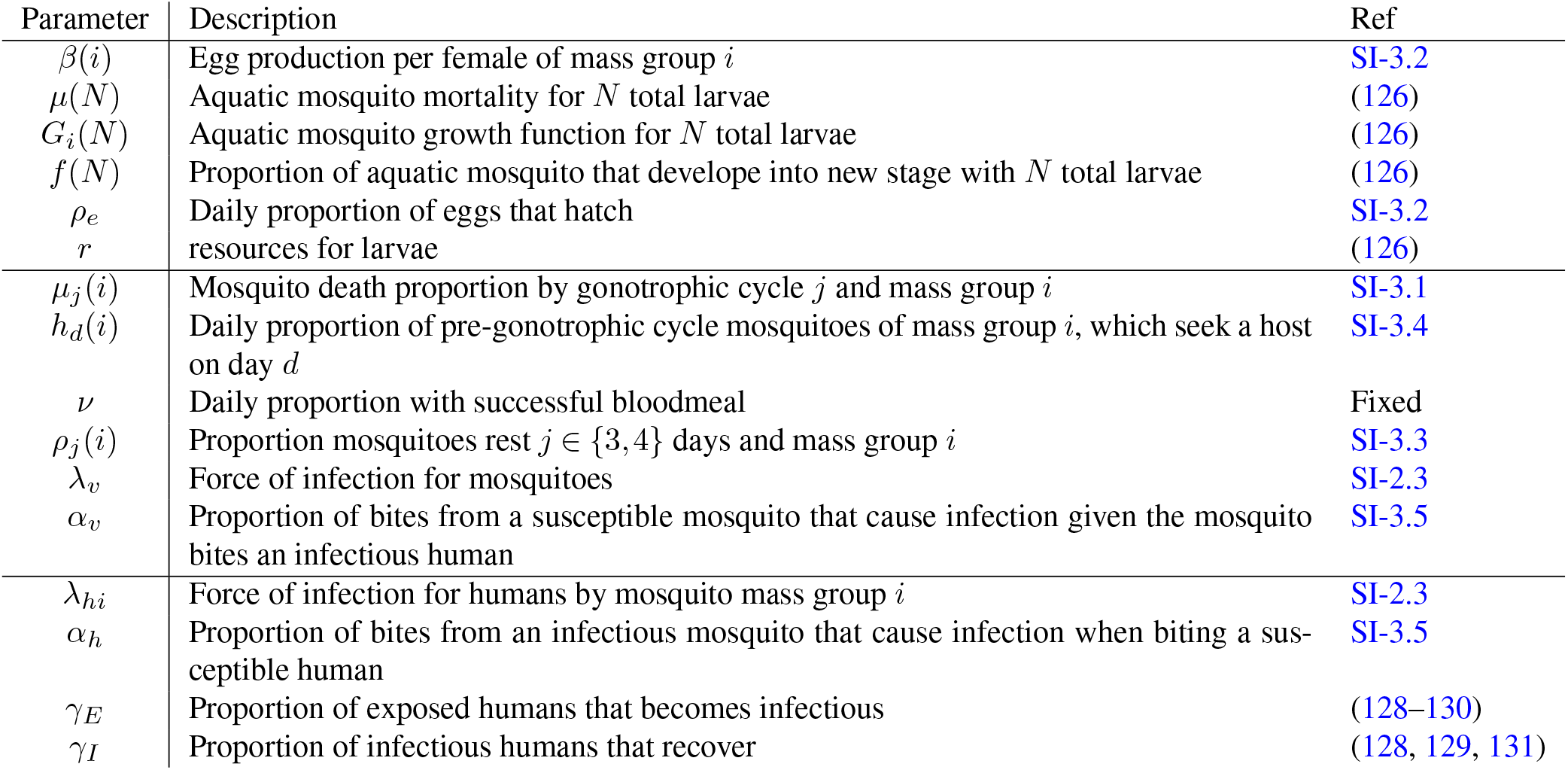
Parameters for LMM.

| Parameter | Description | Ref |
| --- | --- | --- |
| $\beta(i)$ | Egg production per female of mass group $i$ | SI-3.2 |
| $\mu(N)$ | Aquatic mosquito mortality for $N$ total larvae | (126) |
| $G_i(N)$ | Aquatic mosquito growth function for $N$ total larvae | (126) |
| $f(N)$ | Proportion of aquatic mosquito that develop into new stage with $N$ total larvae | (126) |
| $\rho_e$ | Daily proportion of eggs that hatch | SI-3.2 |
| $r$ | resources for larvae | (126) |
| $\mu_j(i)$ | Mosquito death proportion by gonotrophic cycle $j$ and mass group $i$ | SI-3.1 |
| $h_d(i)$ | Daily proportion of pre-gonotrophic cycle mosquitoes of mass group $i$, which seek a host on day $d$ | SI-3.4 |
| $\nu$ | Daily proportion with successful bloodmeal | Fixed |
| $\rho_j(i)$ | Proportion mosquitoes rest $j \in \{3, 4\}$ days and mass group $i$ | SI-3.3 |
| $\lambda_v$ | Force of infection for mosquitoes | SI-2.3 |
| $\alpha_v$ | Proportion of bites from a susceptible mosquito that cause infection given the mosquito bites an infectious human | SI-3.5 |
| $\lambda_{hi}$ | Force of infection for humans by mosquito mass group $i$ | SI-2.3 |
| $\alpha_h$ | Proportion of bites from an infectious mosquito that cause infection when biting a susceptible human | SI-3.5 |
| $\gamma_E$ | Proportion of exposed humans that becomes infectious | (128–130) |
| $\gamma_I$ | Proportion of infectious humans that recover | (128, 129, 131) |

**Table S3.** Variables in RMM.

| Variable | Description |
| --- | --- |
| $N_h$ | Total humans |
| $S_{hj}$ | Susceptible humans of type $j$ |
| $E_{hj}$ | Exposed (infected but not infectious) humans of type $j$ |
| $I_{hj}$ | Infectious humans of type $j$ |
| $R_{hj}$ | Recovered humans of type $j$ |
| $S_v$ | Susceptible mosquitoes |
| $E_v$ | Exposed (infected but not infectious) mosquitoes |
| $I_v$ | Infectious mosquitoes |

**Table S4.**
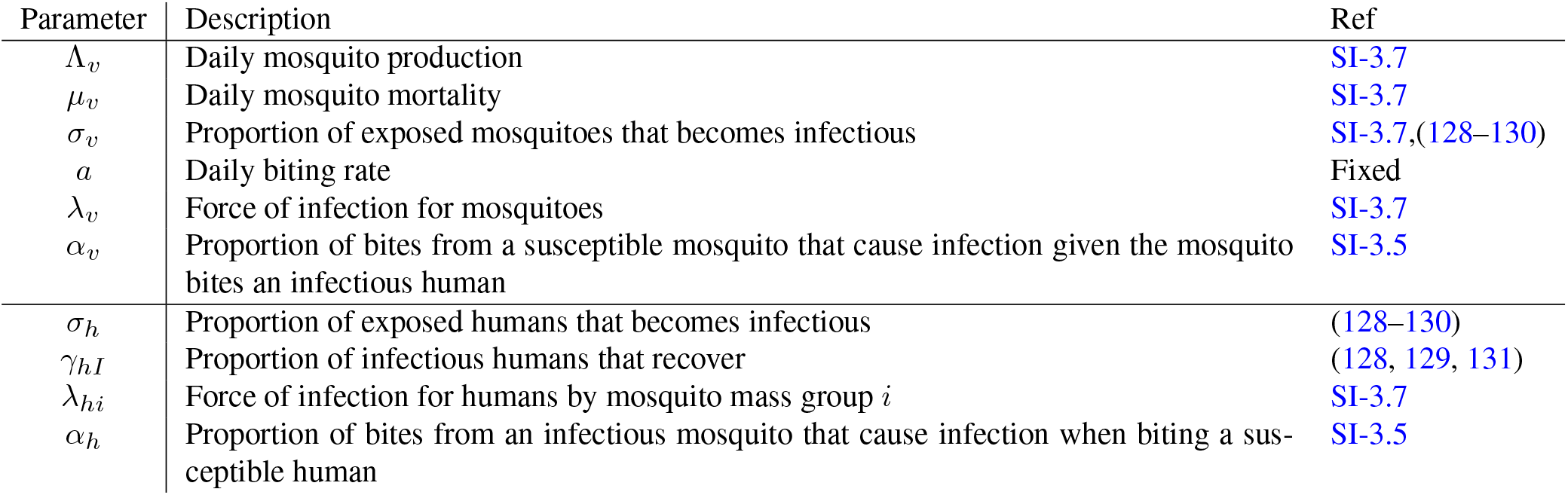
Parameters for RMM.

| Parameter | Description | Ref |
| --- | --- | --- |
| $\Lambda_v$ | Daily mosquito production | SI-3.7 |
| $\mu_v$ | Daily mosquito mortality | SI-3.7 |
| $\sigma_v$ | Proportion of exposed mosquitoes that becomes infectious | SI-3.7,(128–130) |
| $a$ | Daily biting rate | Fixed |
| $\lambda_v$ | Force of infection for mosquitoes | SI-3.7 |
| $\alpha_v$ | Proportion of bites from a susceptible mosquito that cause infection given the mosquito bites an infectious human | SI-3.5 |
| $\sigma_h$ | Proportion of exposed humans that becomes infectious | (128–130) |
| $\gamma_{hI}$ | Proportion of infectious humans that recover | (128, 129, 131) |
| $\lambda_{hi}$ | Force of infection for humans by mosquito mass group $i$ | SI-3.7 |
| $\alpha_h$ | Proportion of bites from an infectious mosquito that cause infection when biting a susceptible human | SI-3.5 |

**Table S5.** Host odor blend composition.

| Compound | Mass (g) | Molar mass (g/mol) | 50X Concentration (M) | Mole fraction ($x_i$) |
| --- | --- | --- | --- | --- |
| Nonanal | 0.01404 | 142.24 | $4.94 \times 10^{-3}$ | 0.4245 |
| Butyric acid | 0.00080 | 88.11 | $4.54 \times 10^{-4}$ | 0.0391 |
| Lactic acid | 0.00841 | 90.08 | $4.67 \times 10^{-3}$ | 0.4014 |
| Octenol | 0.00403 | 128.21 | $1.57 \times 10^{-3}$ | 0.1350 |

**Table S6.**
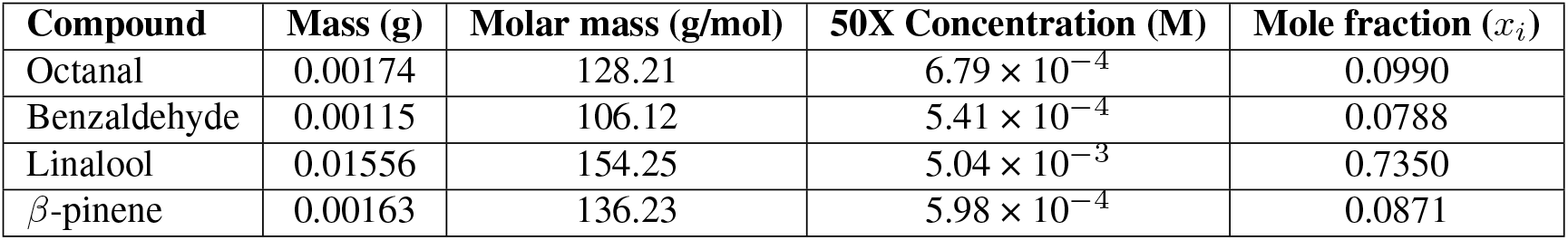
Plant odor blend composition.

## Notes

### Competing Interest Statement

The authors have declared no competing interest.

